# Purchases dominate the carbon footprint of research laboratories

**DOI:** 10.1101/2023.04.04.535626

**Authors:** Marianne De Paepe, Laurent Jeanneau, Jerôme Mariette, Olivier Aumont, Andŕe Estevez-Torres

## Abstract

Despite increasing interest for the carbon footprint of higher education institutions, little is known about the carbon footprint associated to research activities. Air travel and attendance to conferences concentrate recent data and debates but purchases have attracted little attention. Here we develop a hybrid method to estimate the greenhouse gas emissions (GHG) associated to research purchases. To do so, we combine macroeconomic databases, research-centered companies footprints and life-cycle assesments to construct a public database of monetary emission factors (EF) for research purchases. We apply it to estimate the purchases emissions of a hundred of research laboratories in France, belonging to the Labos 1point5 network and gathering more than 20000 staff, from all disciplines. We find that purchases dominate laboratory emissions, accounting for more than 50% of emissions, with a median of 2.7 t CO_2_e/pers, which is 3 to 4-fold the separate contribution from travel, commutes and heating. Median electricity emissions are 5-fold lower in our dataset of laboratories using low carbon electricity but they become preponderant for high carbon electricity mixes (3.5 t CO_2_e/pers). Purchases emissions are very heterogeneous among laboratories and are linearly correlated with budget, with an average carbon intensity of 0.31 *±* 0.07 kg CO_2_e/€ and differences between research domains. Finally, we quantify the effect of a series of demand-driven mitigation strategies obtaining up to *−*20 % in total emissions (*−*40 % in purchases emissions), suggesting that effectively reducing the carbon footprint of research activities calls for systemic changes.

**Significance statement:** Research activities are recently interrogating their contribution to global warming, mainly through the impact of air travel but neglecting the emissions embodied in scientific purchases. However, goods and services used in a research laboratory emit greenhouse gases when they are produced. Here we construct a public and robust database of emission factors to quantify purchases emissions in a laboratory and we use it to assess emissions from a hundred of laboratories in France, from all disciplines. We find that purchases emissions represent half of the of the 6.3 t CO_2_e/pers per year emitted on average per laboratory. Emissions, however, vary greatly between laboratories and disciplines and an analysis of mitigation strategies shows that decreasing demand may significantly reduce purchases emissions.

## Introduction

Planetary boundaries refer to the ensemble of physical, ecological and social constraints that limit the flux of matter and energy sustaining human societies.^1^ They have been a subject of continuous discussion for at least two centuries.^2–8^ This has spurred the necessity for implementing a material accountability, complementary to a monetary one, in order to curb material and energy flows associated to human activities.

Universities and research laboratories have greatly contributed to a better understanding of these planetary limits, in particular concerning global warming^9^ and biodiversity loss.^10^ However, research itself has undesired impacts, both directly by consuming natural resources and generating waste and greenhouse gases (GHG) ^11^ and indirectly through the discovery of processes and techniques that may increase the overall impact of humanity on the environment in the long run.^12–14^

Awareness of the direct impacts of academic research on the environment, and more specifically, on global warming, is illustrated by the steady increase in the scientific literature on the carbon footprint of academic research and higher education. ^15^ In order to quantify GHG emissions in research, two main approaches have been followed: a top-down and a bottom-up approach. In the former, the carbon footprint of entire universities was estimated using aggregated data, in general without distinguishing research and educational activities.^15–18^ In the latter, the footprint of individual and specific research activities such as attending conferences or a PhD project,^19^ scientific events such as international conferences^20^ or disciplines,^21, 22^ were assessed.

The large majority of the footprints estimated by higher education institutions focuses on direct and energy-related emissions^15, 18^ (scope 1 and 2^23^) and only partially includes scope 3 emissions,^24^ i.e. those resulting from activities that occur in locations that are not owned by the institution. They are the most diverse and, therefore, the most difficult to assess, which explains why they are rarely accounted for. Yet, scope 3 emissions, and among them, purchases of goods and services, can represent a large share of their total footprint.^16, 25, 26^ Some studies suggest that they may account for as much as 80% of total emissions.^17, 27^

In this work, we have taken an intermediate approach and selected the research laboratory as a valuable perimeter to evaluate the carbon footprint of research activities. In the first part, we propose a method to estimate the carbon footprint of all the goods and services purchased in the laboratory. We constructed a public listing of monetary emission factors (EFs) associated to 1431 categories of scientific purchases and 61 physical emission factors associated to 8 labware categories using different databases and complementary methods to assess the robustness of our approach. These EFs can be used as is or through the web interface GES 1point5,^28^ an open source free tool for any research laboratory to estimate its carbon footprint. GES 1point5 is developed by Labos 1point5, a nation-wide initiative, launched in 2019, and engaged in a cross-disciplinary estimation of the environmental footprint of research together with the analysis of mitigation strategies. Gathering more than 700 laboratories and more than 300 consolidated GHG inventories, it is possibly the largest database of laboratory emissions worldwide. In the second part of the paper, we analyse 167 GHG inventories associated to 108 distinct French laboratories from all disciplines and show that purchases represent 50% of median emissions. Emissions in general and purchases emissions in particular are very heterogeneous between laboratories and research domains. Interestingly, we find a strong linear correlation between purchases emissions and budget with a carbon intensity of *∼* 0.3 kg CO_2_e/ € for sciences and technology and life and health sciences laboratories and *∼* 0.2 kg CO_2_e/ € for human and social sciences laboratories. We conclude by discussing potential mitigation strategies, showing that reducing purchase-associated emissions is possible but requires systemic changes.

## Results and discussion

Emissions embodied in goods and services can be estimated by measuring physical or monetary flows. To make the problem tractable considering the large number of purchase types in research laboratories, goods were classified according to the French system for accountability in research (NACRES), to which we manually associated monetary emission factors (EFs) in kg CO_2_e/€. Throughout the text all € values correspond to year 2019. The emissions of good *i* were calculated as *e*(*i*) = *p*(*i*) *× EF* (*i*), with *p*(*i*) its price in €. EFs were estimated using the three approaches sketched in Fig. 1: i) an environmentally extended input-output (EEIO) method^29^ that we will call in the following *macro* and note the resulting EFs EF*_macro_*; ii) a process-based method that we will call in the following *micro* (EF*_micro_*); and iii) an intermediate approach based on the carbon footprint of selected companies of the research sector, that we will call in the following *meso* (EF*_meso_*).

**Figure 1:**
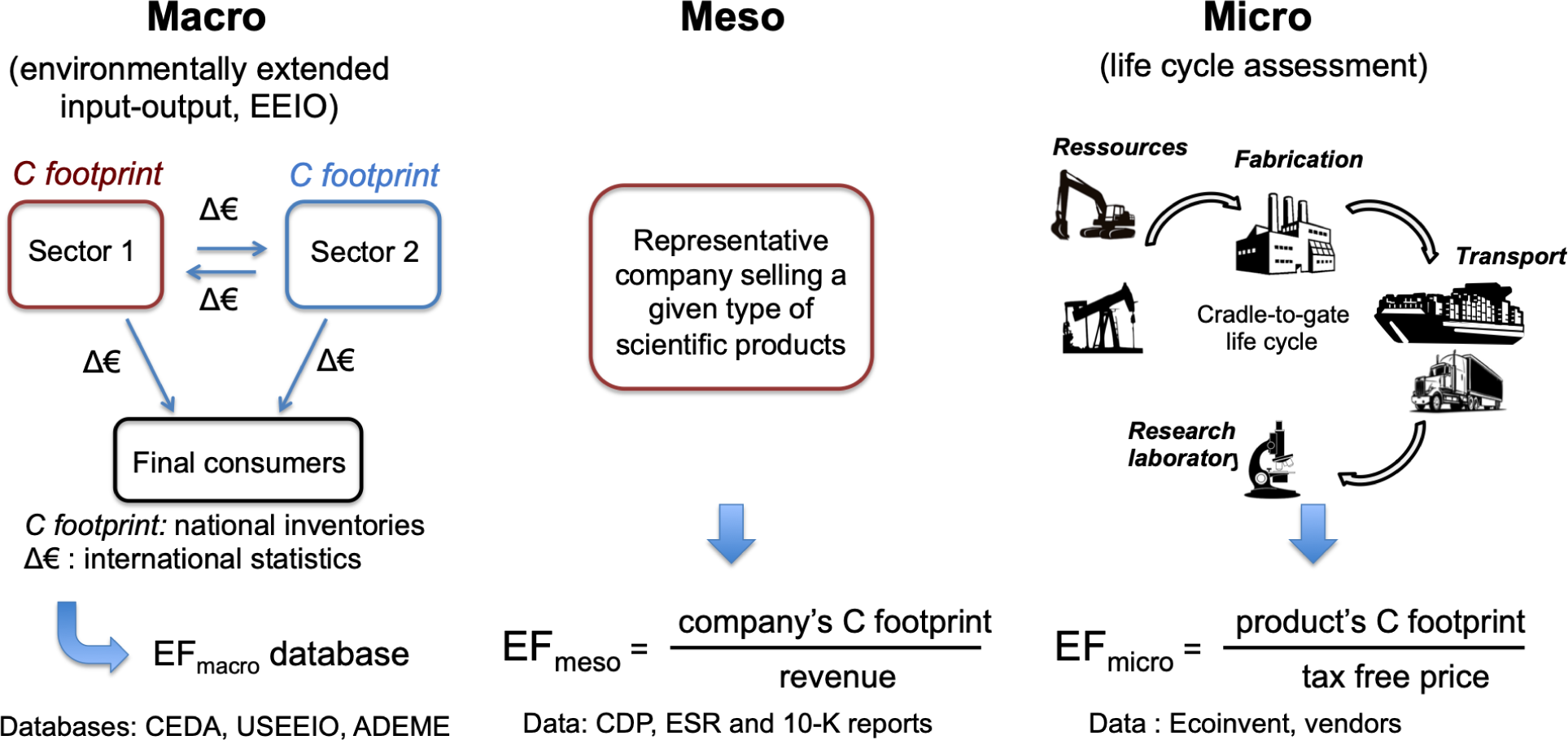
Scheme showing the three approaches used in this work to estimate monetary emission factors (EF) of purchased goods and services.

Environmentally extended input-output (EEIO) methods associate environmental impacts to macroeconomic monetary flows between production and consumption sectors in a given economy or territory.^29^ They have proved useful to estimate the carbon footprint of purchases in large organizations.^30^ However, they should be used with caution when applied to niche products which are abundant in research laboratories. We therefore used a hybrid approach: for purchase categories most specific to research labs (scientific instruments and consumables), we completed the EEIO method by the meso and micro approaches.

### Construction of the emission factor database

In a first step, each of the 1431 NACRES categories identifying goods and services was attributed one or several EFs from each one of three EEIO databases: the two American CEDA^31^ and USEEIO^32, 33^ databases, and the French ADEME^34^ database, the first two providing 430 EFs and the last one 38. This constituted three databases of NACRES monetary EFs, called in the following CEDA, USEEIO and ADEME, respectively. Note that ADEME and CEDA EFs are cradle-to-gate, meaning that transport of goods from the producer to the final consumer are not considered, while USEEIO EFs are cradle-to-shelf, i.e. transportation from the producer to the point-of-sale is included. In a second step, the PER1p5 macro database was constructed by averaging, for each NACRES category, the EFs from the three other databases (Tab. S1). Fig. 2A and Tab. S6 show the properties of the distribution of EFs associated to the different NACRES categories for the four macro databases. Lower EFs are more frequent in the USEEIO database, then comes the CEDA and then the ADEME database with respectively medians of 0.19, 0.27 and 0.40 kg CO_2_e/€. The PER1p5 macro database displays a mean EF that is indeed the average of the means of the other three, with a distribution very similar to the CEDA one although without the very high values (Fig. S1). PER1p5 stands for Purchases emissions in research 1point5.

**Figure 2:**
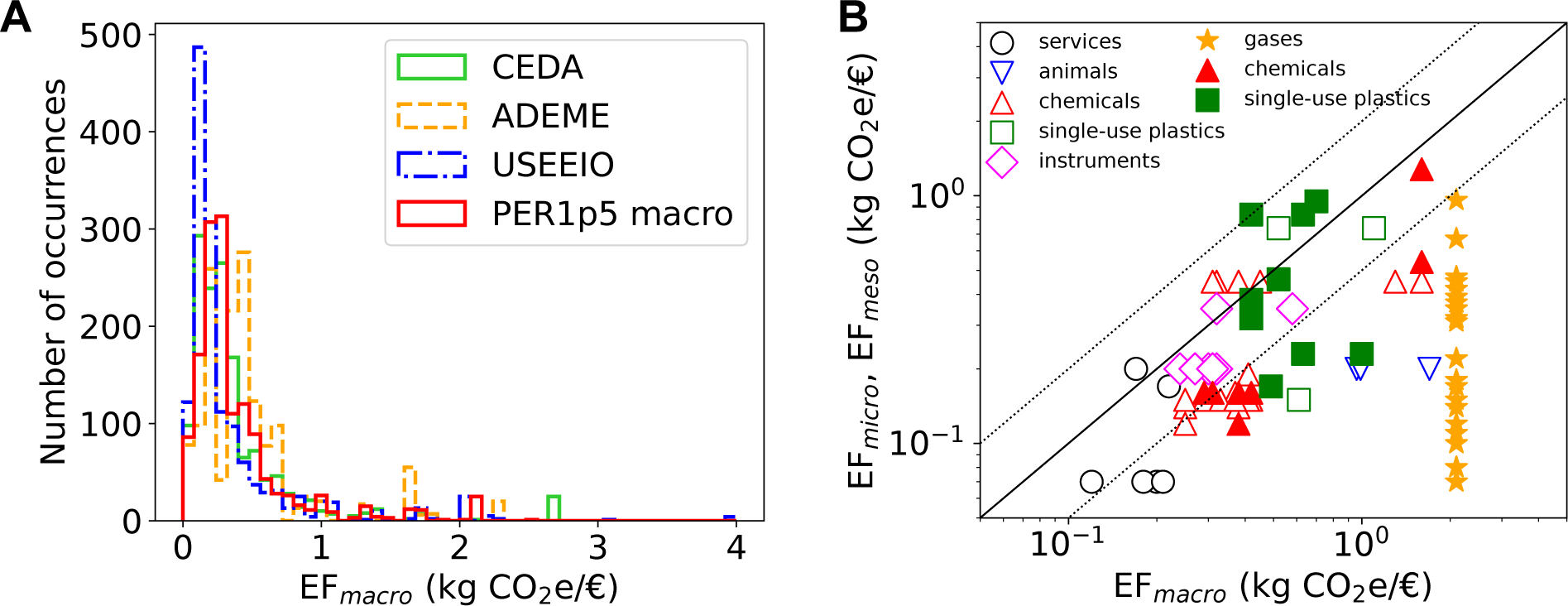
Construction of the PER1p5 NACRES-EF database for estimating emissions associated to purchases in research laboratories. A) Distribution of macro emission factors within the four macro NACRES-EF databases considered in this work. The *y* axis represents the number of NACRES codes assigned to a given EF among the 1431 NACRES codes within the purchases module in GES 1point5. B) Meso (open symbols) and micro (filled symbols) emission factors vs. PER1p5 macro EF for different types of purchases. The plain line indicates identity while the dotted lines refer to 2-fold differences. For readability error bars are shown in Fig. S4.

In a third and final step, the PER1p5 macro database was refined by substituting some macro EFs by meso or micro EFs. Similarly to USEEIO factors, meso EFs cover cradle-to-shelf emissions: they include all upstream emissions (direct and indirect emissions related to the production of goods or services) and emissions of downstream transportation of companies providing instruments, consumables and/or services to the distributors (Tabs. 1 and S2 and S4). As it happens for corporate emissions in other industrial sectors, companies EF*_meso_* most heavily depend on the emissions related to purchased goods and services, that represent 41 to 80% of their total emissions (Tab. S2). 13 EF*_meso_* were determined and attributed to 102 NACRES categories (Tab. S1), with a median of 0.2 kg CO_2_e/€, which is close to the median EF of the USEEIO database. Micro EFs were computed for 60 mono-material products, mostly disposable plastic labware and gas cylinders (Tab. S3) and averaged by NACRES category to obtain 36 EF*_micro_*. These factors were determined by single-impact life cycle assessments^35^ (LCA) that cover the production and transportation to the local supplier. In both meso and micro approaches, the emissions associated with transportation were generally below 10%, except for some gas cylinders (up to 40% for high purity nitrogen) and some plastics and chemicals (*<* 16%) (Tabs. S2-S3).

Fig. 2B shows the correlation between micro/meso EFs and macro ones. Differences between the EFs are expected as each approach suffers from truncation and aggregation issues.^37^ In the macro EEIO factors, capital goods are not considered, which tends to underestimate EF of commodities necessitating important material investments. By contrast, all activities not directly included in manufacturing activities (business travel, employee commuting, waste, purchases of other goods and services and non-attributable processes) are not considered in LCA. Here we therefore performed LCA only for goods with very simple production procedures, and attempted to limit this truncation (see SI methods). Finally, the precision of our meso EFs, though not suffering from any truncation issue, is limited by an important aggregation issue, as companies do not produce a single type of product. Yet, for a given category, on average, EF*_meso_* are of the same order of magnitude than EF*_macro_*. Exceptions concern companies producing chemicals and animals for research, commodities which are not well represented in EEIO databases due to aggregation with very different products (chemicals for industry and livestock breeding). For categories corresponding to single-use plastics, with a single exception, EF*_micro_* were close to EF*_macro_* (less than a 2-fold difference). However, EF*_micro_* were much lower than EF*_macro_* for chemicals, laboratory glassware and especially gas cylinders. Here again, this most probably reflects aggregation issues in EEIO categories, as gases are bought by laboratories in much smaller volumes compared to industries, resulting in much higher prices per kg of gas. With some exceptions (see Methods), these micro and meso EFs substituted the corresponding macro EFs in the PER1p5 macro database to constitute the final PER1p5 database. 9 % of EFs were changed (7 % with meso EFs and 2% with micro EFs, Tab. S1 and Figs. S2-S3). We assigned an 80 % relative uncertainty to each EF as it is common practice for monetary EFs (see Methods).

### The distribution of carbon intensities in the laboratory research economy

A French research laboratory is an administrative and scientific unit that typically gathers 50-400 staff, including researchers, professors, technicians and administrators. It can be assimilated to a university department. To gather financial data from public French laboratories to estimate their purchases emissions we relied on GES 1point5, ^28, 38^ an online, free, open source tool developed by the Labos 1point5 research project.^39^ We created a purchases module that allowed volunteer laboratories to upload their expenses associated to NACRES purchases categories. Interestingly, GES 1point5 allows laboratories to estimate other emission sources such as scope 1 (owned vehicles, cooling gases), scope 2 (electricity and heating) and scope 3 (travels, commuting and computer devices) associated emissions. We designed the purchases module to avoid double counting with the emissions taken into consideration by the other modules. Note that for methodological reasons we have access to the cost of goods and services declared through the purchases module but not for those declared through the computer devices one. For this reason, in the following we will use the term *purchases* to refer to the emissions of all goods and services declared through both modules (as in Figs. 4 and 6) and the term *purchases module* for emissions associated to this module alone, in particular when we need to correlate purchases emissions and budget (as in Figs. 3 and 5). Among the 750 research laboratories in the Labos 1point5 network, 108 laboratories submitted 167 GHG purchases inventories for different years (mostly 2019-2021, Fig. S5). When suitable (Figs. 3 and 4) submissions from different years were averaged for each lab. This averaging only marginally changed the results (Fig. S7).

**Figure 3:**
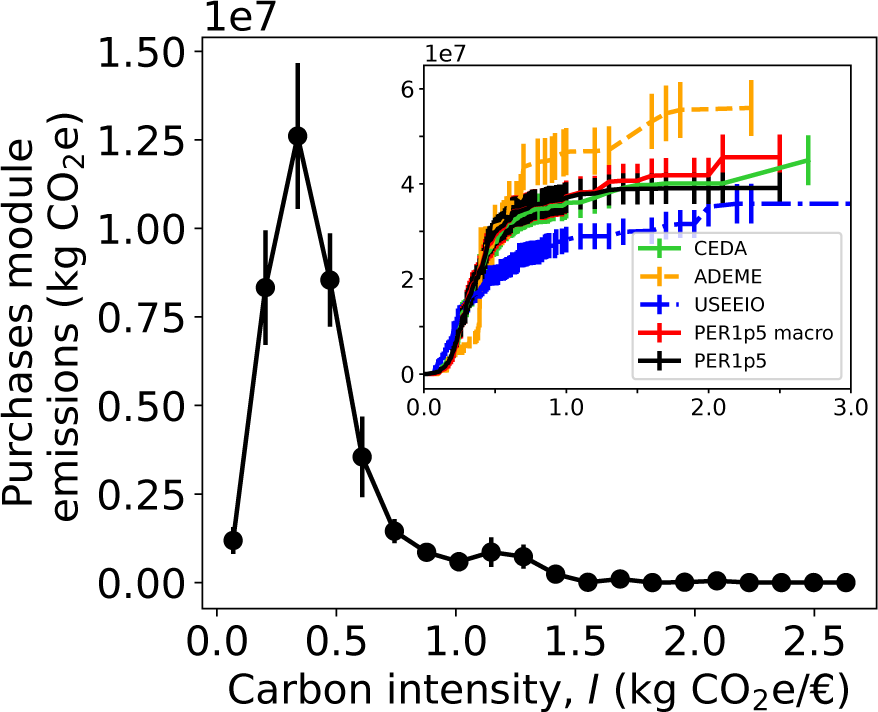
Distribution of carbon intensities within the GES 1point5 laboratory emission database calculated with PER1p5 EFs. For each bin in the *x* axis, the corresponding carbon intensity *I* is multiplied by the total amount of purchases in € to calculate the purchases emissions associated to that *I*. The inset shows the cumulated distribution of carbon intensities for all NACRES-EF databases. *n_s_* = 167 GHG submissions averaged over *n_l_*= 108 distinct laboratories.

**Figure 4:**
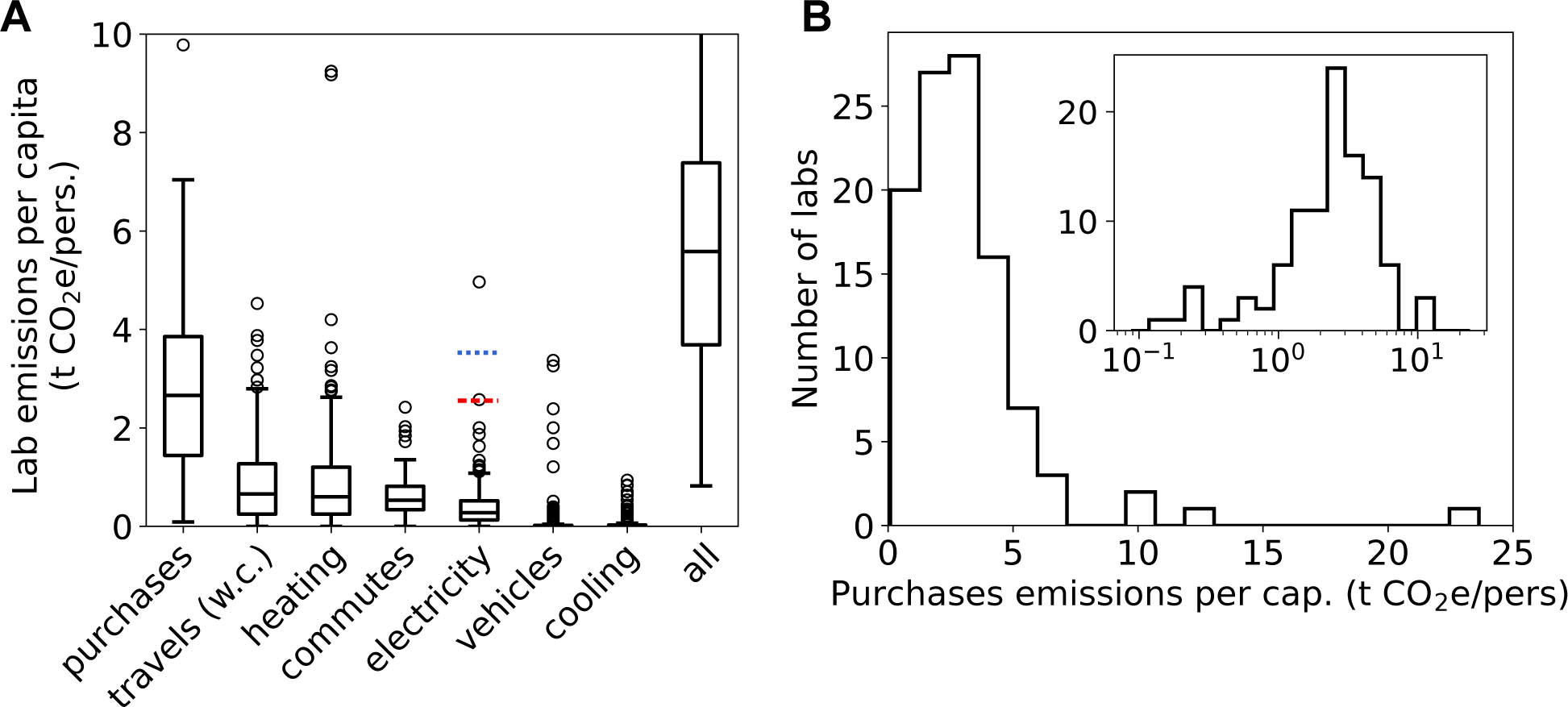
Purchases dominate GHG emissions among laboratories using low-carbon electricity. A) Boxplot of laboratory emissions per capita per emission source. *n_l_ ≥* 190 for all types except for purchases (*n_l_* = 105). w.c. indicates that emissions associated to plane transportation were calculated with contrails.^28^ Electricity emissions are calculated for three different mixes: French mix (boxplot in black), world mix (median as a dashed red line), and high-carbon mix (median as dotted blue line). Note that the *y* axis is truncated (see Fig. S8 and panel B). B) Distribution of purchases emissions per capita, the inset shows the same data in log scale. Purchases emissions calculated with the PER1p5 NACRES-EF database.

**Figure 5:**
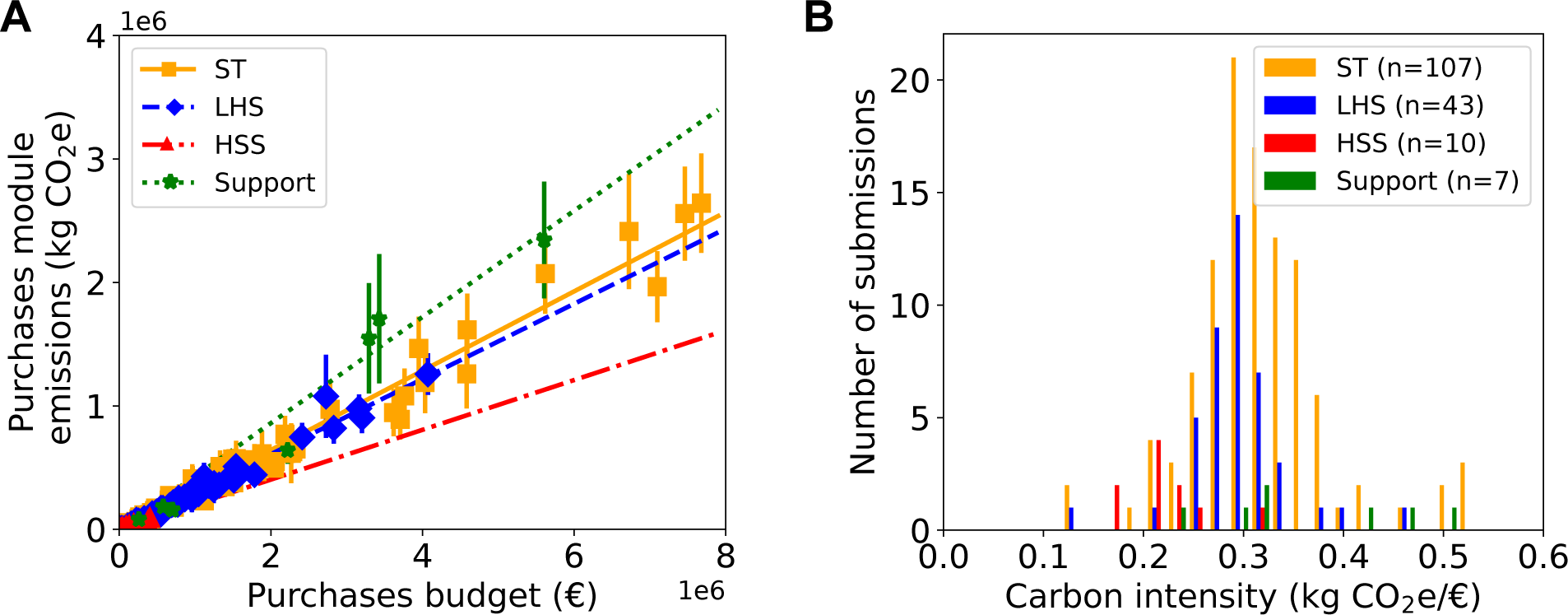
Purchases emissions are proportional to budget, with differences between research domains. A) Purchases module emissions vs. budget for all GHG laboratory footprints in the GES 1point5 lab emission database. Error bars corresponds to one standard deviation calculated as described in Methods. Lines are linear fits with zero intercept, whose results are provided in Tab. S11. B) Histogram of purchases module carbon intensities for different scientific domains. HSS: Human and social sciences, LHS: Life and health sciences, ST: Science and technology. *n_s_* = 167 GHG submissions associated to *n_l_* = 108 laboratories.

**Figure 6:**
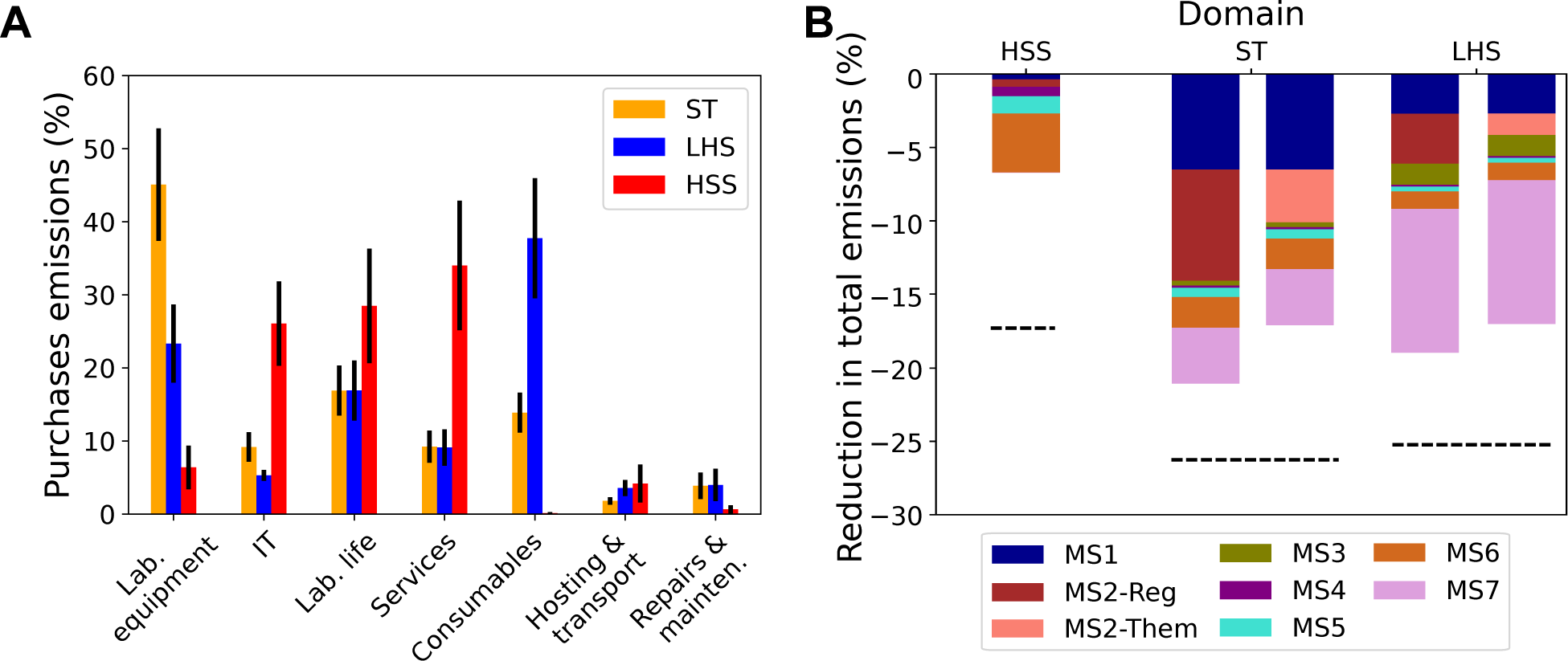
Typology of purchases emissions and quantification of mitigation strategies. A) Share of purchases emissions per research domain (colors) broken down by purchases category. Error bars correspond to one standard deviation and letters indicate significant differences (*p <* 0.05). *n_s_* = 162 purchases submissions averaged over *n_l_* = 105 laboratories. B) Relative reduction of the total laboratory emissions by research domain expected within the GES 1point5 lab emission database for the seven mitigation strategies considered. MS1: +50% of lab equipment life-time; MS2: 50% pooling of lab equipment, either by region (-Reg) or by research sub-discipline (-Them); MS3: replace 80% of plastic by glass; MS4: 75% vegetarian catering; MS5: *−*50% in furniture purchases; MS6: *−*50% in IT purchases; MS7: *−*50% in consumable purchases. Dashed lines correspond to *−*50% in purchases emissions. *n_s_* = 135 submissions corresponding to *n_l_* = 93 laboratoires. ST: science and technology (*n_l_* = 64), LHS: life and health sciences (*n_l_* = 23), HSS: human and social sciences (*n_l_* = 6) laboratories.

Figs. 3 and S6 show the distribution of carbon intensities *I* in the ‘research laboratory economy’ captured by our data. Note that we use the term carbon intensity for average EFs while we use EF for a given good. Carbon intensities are weighted by the associated purchases emissions from all laboratories calculated for the five NACRES-EF databases considered here. The distribution of intensities with PER1p5 displays two peaks, a large one at 0.34 kg CO_2_e/€ and a smaller one at 1.2 kg CO_2_e/€, accounting respectively for 60% and 17% of emissions, and yielding an average intensity 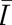= 0.31 kg CO_2_e/€ (Tab. S6). CEDA and PER1p5 macro provide similar distributions for *I <* 1.0 but with larger emissions at higher intensities, resulting in a larger 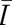 of 0.34 and 0.35 kg CO_2_e/€ respectively. USEEIO and ADEME provide extreme distributions with the former attributing lower emissions in the range 0.2 *−* 0.6 kg CO_2_e/€ and higher emissions for *I >* 1.5, which yields 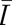 = 0.28 kg CO_2_e/€, and the later displaying significantly larger emissions for *I >* 0.7 kg CO_2_e/€, associated with a higher average (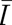 = 0.43 kg CO_2_e/€). Goods associated to EF*_micro_* and EF*_meso_* in PER1p5 represented 12% of purchases expenses on average, with high disparity from one lab to another (from 0 to 53%). As a result, cumulated purchases emissions in the database dropped by 16% when PER1p5 instead of PER1p5 macro was used (Tab.S7).

These results highlight the interest of using different NACRES-EF databases to estimate purchases emissions as we can evaluate, at least partially, the uncertainties of the results. From the different NACRES-EF databases (Fig. 3 inset), we conclude that the average carbon intensity of laboratory purchases is in the range 0.22 *−* 0.42 kg CO_2_e/€, i.e. 0.32 *±* 0.10 kg CO_2_e/€, the lower value given by USEEIO and the larger one by ADEME. This implies that the purchases emissions aggregated for all laboratories could be estimated with a precision of 30 % by just multiplying the purchases budget by this average carbon intensity.

### Purchases and electricity dominate laboratory emissions

We now have a robust method to estimate laboratory purchases emissions and in the following we will use solely PER1p5 EFs to calculate them. An important question is the relative importance of each emission source as this conditions where the efforts of reduction need to be concentrated. Fig. 4A and Tab. S8 display the distribution of emissions for the seven types of emission sources in the GES 1point5 lab emission database. Importantly, this perimeter includes all upstream and in-house laboratory emissions except those due to investments not carried by the laboratories themselves but by the hosting institutions (such as construction and large scientific infrastructures), waste, staff meals and some specific direct emissions (see Methods). This database contains more than 300 GHG emission inventories from more than 200 laboratories employing more that 40000 staff, except for purchases for which more than 160 inventories from more than 100 different laboratories and employing more than 23000 staff were available. Within the considered perimeter, average laboratory emissions add up to 6.2 t CO_2_e/pers. (median 5.6), with purchases accounting for *∼* 50 % of the share and a median of 2.7 t CO_2_e/pers. Travels, heating and commuting to work are far weaker with 10-15% and a median of 0.5-0.7 t CO_2_e/pers. Electricity (8%, 0.3 t CO_2_e/pers.) comes next, with electricity being particularly low in our dataset due to the low carbon emissions of the French electricity system (60 g CO_2_e/kWh^40^). Emissions associated to lab-owned vehicles and cooling systems are negligible on average. Laboratory emissions are however very heterogeneous and the distributions of per capita emissions per source are wide. In particular, for purchases emissions quartiles were (1.5, 3.8) t CO_2_e/pers and extreme values spanned more than two orders of magnitude (0.09 *−* 29 t CO_2_ e/pers, Fig. 4B).

To compare these data internationally we corrected by the carbon intensity of the electricity mix used in the laboratory. The average carbon intensity of the world electricity mix is 7.9-fold higher (475 g CO_2_e/kWh^41^), while the highest electricity intensities can be up to 11.7-fold higher (700 g CO_2_e/kWh^42^). In these cases the median of electricity emissions would equal the median of purchases emissions per capita (2.4 t CO_2_e/pers) or become preponderant (3.5 t CO_2_e/pers).

### Purchases emissions are correlated to budget and research domain

Figs. 5 and S12 show that purchases emissions are linearly correlated to purchases budget with variations by research domain. In contrast, the correlation between emissions and number of staff was weaker (Fig. S10). Laboratory budgets in our database spanned 2 *×* 10^3^ *−* 8 *×* 10^6^ € with a symmetric distribution of carbon intensities of mean 0.31 kg CO_2_e/€ and a s.d. of 0.07 CO_2_e/€ very close to the carbon intensity resulting from linear fitting (0.33 kg CO_2_e/€) (Fig. S11 and Tab. S11). Human and social sciences (HSS) laboratories displayed significantly lower carbon intensities (0.22 kg CO_2_e/€) while support laboratories, i.e. large experimental platforms that provide analysis services, display larger carbon intensities associated to a wider distribution (0.43 kg CO_2_e/€, Tab. S11). Science and technology (ST) and life and health science (LHS) laboratories were associated to carbon intensities close to the mean (0.32 and 0.30 kg CO_2_e/€, respectively).

### The typology of purchases emissions depend on research domain

We classified purchases into seven categories: consumables, IT, lab equipment, repairs & maintenance, services, transport & hosting not included in travel and commuting, and laboratory life (see SI Methods). The share of emissions for these categories strongly depended on the research domain (Fig. 6A). ST purchases emissions are dominated by the acquisition of laboratory equipment (44 *±* 8 %), while for LHS consumables dominate (38 *±* 8 %). HSS exhibit a clearly different typology with three categories with shares close to 30% of emissions: services, laboratory life and IT. Weaker but still important contributions for ST laboratories are laboratory life, IT, consumables and services, while for LHS laboratories these are equipment, laboratory life, IT and services. Emissions associated to hosting during travels and to repairs and maintenance represent 5% or less of the purchases footprint for the three domains. Table 2 sumarizes the average EFs per category and domain. These factors can be easily used by any laboratory to estimate their purchases emissions.

**Table 1:** Meso emission factors (corporate direct and upstream emissions divided by total sales), in kg CO_2_e/€, of companies whose main clients are research laboratories, aggregated by business segment. Details by company are given in Tabs. S2 and S4. Data calculated from corporate reports and ref. 36.

| Business segment | EF |
| --- | --- |
| Gloves and hygienic equipment | 0.74 |
| Chemicals | 0.45 |
| Global lab supplier (Instrumentation, consumables & services) | 0.13 – 0.38 |
| Scientific equipment ($> 80\%$ of sales) | 0.18 – 0.35 |
| Biotech consumables | 0.14 – 0.16 |
| Scientific services | 0.07 – 0.19 |

**Table 2:** Average emission factors (in kg CO_2_e/€) for purchases module categories for different domains: Human and social sciences (HSS), life and health sciences (LHS), sciences and technology (ST) and an average of all three (and thus excluding Support labs).

| Category | HSS | LHS | ST | ALL |
| --- | --- | --- | --- | --- |
| Consumables | 0.94 | 0.37 | 0.51 | 0.44 |
| Hosting - transport | 0.30 | 0.43 | 0.45 | 0.44 |
| IT w/o devices | 0.30 | 0.27 | 0.20 | 0.21 |
| Lab. equipment | 0.31 | 0.32 | 0.34 | 0.33 |
| Lab. life | 0.36 | 0.46 | 0.38 | 0.39 |
| Repairs - maintenance | 0.21 | 0.22 | 0.23 | 0.23 |
| Services | 0.14 | 0.13 | 0.15 | 0.14 |

The differences displayed in Fig. 6A imply that mitigation strategies should consider the scientific specificity of the laboratories. At the scale of a single laboratory, our method allows a finer view of the distribution of emissions among different purchases subcategories (Fig. S14). However, one must keep in mind that the financial categorization used here to identify purchases (NACRES) does not allow to distinguish between similar goods with potentially different carbon footprints, thus jeopardizing the estimation of supply-driven mitigation strategies, i.e. decreasing the emission factors.

### Identifying and quantifying mitigation strategies for scientific purchases

Despite these limitations, it is possible to evaluate the effect of demand-driven mitigation strategies that involve reducing the purchase of certain items. We considered seven of such strategies applied to the three scientific domains (Fig. 6B) and we quantified their relative effect compared to the total emissions of the laboratory (and not just purchases emissions). Two mitigation strategies addressed scientific equipment: a 50% increase in equipment service life (MS1) and the pooling of 50% of equipments either by sub-discipline (MS2-Them) or by region (MS2-Reg). Two strategies focused on laboratory-life purchases: a 75% conversion of laboratory-paid catering to vegetarianism (MS4) and a 2-fold reduction in furniture purchases (MS5). Two strategies concerned consumables: replacing 80% of plastic consumables by glass (MS3) and reducing 50% all consumables purchases (MS7). Finally, we considered the effect of reducing by 50% IT purchases (MS6). Note that our analysis neglects potential rebound effects that may increase electricity or commuting emissions if these strategies were implemented. As expected from Fig. 6A, the impact of these strategies was relatively similar for ST and LHS laboratories and different for HSS ones. For ST, the most effective strategies concerned pooling equipment by region (MS2-Reg), increasing equipment life-time (MS1), reducing consumables (MS7), and reducing IT (MS6). For LHS, MS7 was clearly the most effective followed by equipment pooling by region (MS2-Reg) and increasing life-time (MS1). Then came replacing plastic by glass (MS3) (our results being in agreement with ref. 43) and reducing IT. Reducing furniture and conversion to vegetarianism was negligible for both domains. For HSS, reducing IT purchases was the most effective, followed by conversion to vegetarianism. The addition of all seven strategies decreased purchases emissions by *∼* 40% and total emissions by *∼* 20%, i.e. 1.3 t CO_2_e/pers. on average, both for ST and LHS laboratories. In contrast, for HSS, the purchases footprint reduction was *∼* 20% and the total one was *∼* 6%, i.e. 0.2 t CO_2_e/pers. on average (Fig. S15). We conclude that demand-driven mitigation strategies may be very effective to reduce emissions of both ST and LHS laboratories.

## Discussion and conclusion

Purchases emissions are almost systematically neglected^15, 18, 25^ when calculating the carbon footprint of higher education institutions, except in few seminal studies.^16, 17, 44^ However, these works do not separate research and teaching activities, they use a single set of monetary EFs and they only analyse a single institution.

Interestingly, the average carbon intensity calculated by Larsen et al. for a Norwegian technical university,^16^ 0.39 kg CO_2_e/€ 2019, is close to the one calculated here for a French database of more than hundred different laboratories (0.31*±*0.07 kg CO_2_e/€ 2019). However, Larsen et al did not find significant differences in the carbon intensities between research domains (Tab. S13), in particular with HSS, in contrast to the current work. We thus hypothesize that this difference results from the separation of research from teaching activities in our work. Such distinction is important as our data suggest that mitigation strategies will need to be adapted to each research domain. However, the results obtained for HSS laboratories need to be considered with caution because at the time of our study only 10 inventories from 8 distinct laboratories were available in the GES 1point5 laboratory emission database (Figs. S16-S17).

In addition, available data of purchases footprints in universities rely on either non-public EF^16^ or exclusively from general-economy EEIO EF databases such as EXIOBASE,^45^ thus not offering specific factors for research laboratories. By comparing three EEIO EF databases and hybridizing them with LCA and company data for selected goods specific to research activities, our work provides laboratories around the World with a database of emission factors to easily calculate purchases emissions with different granularities, either using EFs in Tabs. S1 or 2, in addition to valuable meso monetary and micro physical EFs in Tabs S2 and S3. Our results suggest that the PER1p5 EF database allows to calculate laboratory purchases emissions with a 30% precision. To improve the precision further work is needed, in particular to refine emissions associated to laboratory instruments. A crude LCA estimate for mass spectrometers^46^ yields 16 tCO_2_e/t of instrument, i.e. *∼* 0.02, kgCO_2_e/€ and a more detailed calculation for a chromatography apparatus^47^ gives 0.6 tCO_2_e/unit, i.e. 0.03 *−* 0.12 kgCO_2_e/€, while in PER1p5 the typical EF for lab equipment is 0.3 kgCO_2_e/€. Indeed, research instruments are manufactured in small series while LCA gives accurate results only for mass-produced products that have high production costs relative to services such as research & development, administrative and commercial costs.^48^ For instance, unique instruments such as satellites production emissions calculated though a hybrid method are 8 times higher than those estimated through LCA.^49^ At this point we think that the PER1p5 monetary estimate is more reliable.

By analysing a unique dataset of GHG inventories from a hundred of laboratories we have shown the great heterogeneity of emissions among research laboratories, both between different emission sources and within purchases alone. Importantly, our data suggest that, within a given research discipline, laboratory budget is linearly related to purchases emissions, in a similar way as income is the main driver of the carbon footprint of households, though in the latter these effects are not linear. ^50^ The strong linearity observed between purchases emissions and budget in Fig. 5A is intriguing. On the one side, one may argue that this linearity is consubstantial to a model using monetary EFs, and thus it is not a result per se. On the other hand, the distribution of carbon intensities in our data (Figs. 3 and 5B) is relatively large, and thus suggests that both the linearity and the differences in the carbon intensities observed between domains are a result and not an artefact of our model.

Finally, our demand-based mitigation analysis highlights that experimental laboratories would effectively reduce emissions by developing strategies to diminish equipment, consumables and IT purchases, in particular by extending their lifetime and through sharing. For human and social sciences purchases represent a smaller share of total emissions and thus their contribution to mitigation is lower, but increasing the lifetime of IT equipment still represents a significant reduction. In addition to these demand-based mitigation strategies, on the long term, the general decarbonation of worldwide industry may induce a decrease in EFs and thus in total emissions.

In summary, our work provides a unique, public and curated database of EFs to estimate purchases emissions in a laboratory, it shows that purchases dominate laboratory emissions and ranks the usefulness of mitigation strategies by research domain.

## Methods

### Classification of goods and approach

Services and goods purchased in a laboratory are classified according to the French NACRES nomenclature, used in the accountability of the majority of research institutions in France. ^51^ There are 1431 defined NACRES codes split into 24 large categories (Tabs. S5 and S1). In this work, each NACRES code is given an EF covering GHG emissions associated to all stages of its production (cradle-to-shelf perimeter). Each NACRES code is given an EF using the *macro* method (see below), and certain types of goods were also attributed a meso or a micro EF (see below), that were used to construct the final hybrid database PER1p5, which contained 1281 macro, 108 meso and 43 micro EFs (Tab. S1). Complete methodology is described in the SI Methods.

### The macro approach

To associate EFs with each NACRES, we used three different EEIO databases of monetary emission factors: the French *Ratios Monétaires* database published by the *Agence De l’Environnement et de la Ma^ıtrise de l’Energie* (ADEME) in 2016; the U.S. CEDA^31^ database provided by Vitalmetrics (version 4.8 released in 2014); and the U.S. Supply Chain GHG Emission Factors for US Commodities and Industries calculated from the USEEIO models^32, 33^ compiled by the US Environmental protection agency (EPA, published in 2018). Both American databases contain approximately the same 430 categories, while the French ADEME database provides monetary factors for only 38 categories.^34^ As the NACRES types cannot always be associated to a single category of the EEIO databases, we associated up to 2 ADEME EFs and up to 6 CEDA/USEEIO EFs to each NACRES category (Tab. S1). We proceeded heuristically by assigning all the EEIO categories of commodities that have similarities (in terms of composition and/or manufacturing process) with the products comprised in each NACRES type. To provide a single EF for each NACRES we averaged the allocated EFs, first within each database, and then between databases to yield the PER1p5 macro database. For each EF a uniform relative uncertainty of 80% was attributed to all EFs.

### The meso approach

To consolidate macro NACRES-EF database, we used a supplier-based approach, using GHG emissions and financial data of companies whose main segments of activity are to manufacture products or provide services to the research, analytical and health markets. We gathered emission data from the Carbon Disclosure Project (CDP)^36^ or from internal reports, and financial data from the annual reports of companies. The emission categories used encompass all upstream activities involved in the production of goods or services, similarly to the cradle-to-gate perimeter of EEIO databases, but also downstream transportation as most shipment costs are included in prices for laboratory products. The meso monetary EFs are then computed as *EF_meso_* = (scope 1+2+3 upstream emissions)*/*(revenue).

### The micro approach

For laboratory mono-material products that represented important purchases from a panel of laboratories, we performed single impact cradle-to-gate LCA. This concerned 60 products distributed in 28 NACRES categories, such as all gases and some plasticware and glassware (Table S3). LCA included raw material manufacturing, item manufacturing and transport to the local supplier. Emission factors of each step were obtained from the Ecoinvent database version 3.8. The product monetary EFs are then computed by dividing the product carbon footprint by its price. The micro monetary EFs are then computed as the mean of the monetary EFs of all products belonging to the same NACRES category (1 to 6 products by NACRES category).

### Data collection and treatment

All data used in this study were collected with the GES 1point5 web application.^28, 38^ Volunteer French research laboratories submitted their purchase data through the purchase module of GES 1poin5 as a csv file with NACRES codes and the associated tax-free purchase price. Since heating, electricity, commuting, professional travels and computers were already included in GES 1point5 as dedicated modules, each NACRES code was associated to a ‘Module’ tag taking five different values: *purchase*, *energy*, *vehicles*, *travel* and *computer* (Tab. S1). The monetary approach described here is only used to calculate the emissions of the NACRES codes labeled *purchase*. Purchases emissions are the sum of emissions calculated via the purchases module (via monetary EFs) and the computer devices module (via physical EFs) of GES 1point5, the former clearly dominating purchases emissions. Emissions related to the other sources were computed differently by the dedicated modules of GES 1point5 with EFs based on physical flows as described in ref. 28. The definition of the research domains is given in SI Methods.

Data analysis was performed using custom Python routines. NACRES codes were classified in 7 categories: *lab.life* (food, landscaping, leisure, building), *consumables* (raw materials, chemicals/biologicals and living organisms), *lab.equipment* (laboratory equipment and instruments), *hosting* (professional travel, including lodging and taxi but excluding all other transport), *info* (computers and audio-video equipment), *services* and *maintenance*. Per capita emissions were calculated by full-time equivalent in research, each staff counting 1 except professors counting 0.5.

### Mitigation strategies

Calculations for the seven mitigation strategies are detailed in SI Methods.

## Acknowledgement

We thank Lucille Zribi, Christophe Brun, Franck Davoine and Aymeric Serazin for early developments of the NACRES-EF database, Tamara Ben-Ari, Anthony Benoist and Philippe Loubet for helpful discussions, Cyril Bernard for help with EXIOBASE, Odile Blanchard, Samuel Calvet, Julian Carrey, Fabienne Gauffre, Etienne-Pascal Journet, Sandrine Laguerre, Arthur Leblois, Anne-Laure Ligozat, Elise Maigne, Gerald Salin and Zhou Xu for testing the purchases module of GES 1point5 and the hundreds of research staff that uploaded GHG inventories to the GES 1point5 laboratory emission database. This work was conducted as part of the research network GDR Labos 1point5, funded by ADEME, CNRS and INRAE. It has also been partially supported by the overheads of the European research council grant Mesomat.

## References

(1) Steffen, W. et al. Planetary boundaries: Guiding human development on a changing planet. Science 2015, 347.

(2) Jevons, W. The Coal Question; Macmillan, 1865.

(3) Georgescu-Roegen, N. The entropy law and the economic process; Harvard University Press, 1971.

(4) Meadows, D. H.; Meadows, D. L.; Randers, J.; Behrens III, W. W. The limits to growth; 1972.

(5) Hansen, J.; Nazarenko, L.; Ruedy, R.; Sato, M.; Willis, J.; Del Genio, A.; Koch, D.; Lacis, A.; Lo, K.; Menon, S.; Novakov, T.; Perlwitz, J.; Russell, G.; Schmidt, G. A.; Tausnev, N. Earth’s Energy Imbalance: Confirmation and Implications. Science 2005, 308, 1431–1435.

(6) IPCC, Climate Change 2014: Synthesis Report. Contribution of Working Groups I, II and III to the Fifth Assessment Report of the Intergovernmental Panel on Climate Change; Report, 2014.

(7) Steffen, W.; Broadgate, W.; Deutsch, L.; Gaffney, O.; Ludwig, C. The trajectory of the Anthropocene: The Great Acceleration. The Anthropocene Review 2015, 2, 81–98.

(8) Jürgen, R. The Evolution of Knowledge: Rethinking Science in the Anthropocene. HoST - Journal of History of Science and Technology 2018, 12, 1–22.

(9) IPCC, Synthesis report of the IPCC sixth assessment report (AR6); Report, 2023.

(10) IPBES, Global assessment report on biodiversity and ecosystem services of the IPBES ; Report, 2019.

(11) Urbina, M. A.; Watts, A. J. R.; Reardon, E. E. Labs Should Cut Plastic Waste Too. Nature 2015, 528, 479–479.

(12) Brockway, P. E.; Sorrell, S.; Semieniuk, G.; Heun, M. K.; Court, V. Energy efficiency and economy-wide rebound effects: A review of the evidence and its implications. Renewable and Sustainable Energy Reviews 2021, 141, 110781.

(13) White, L. The Historical Roots of Our Ecologic Crisis. Science 1967, 155, 1203–1207.

(14) Voulvoulis, N.; Burgman, M. A. The Contrasting Roles of Science and Technology in Environmental Challenges. Critical Reviews in Environmental Science and Technology 2019, 49, 1079–1106.

(15) Valls-Val, K.; Bovea, M. D. Carbon footprint in Higher Education Institutions: a literature review and prospects for future research. Clean Technologies and Environmental Policy 2021, 23, 2523–2542.

(16) Larsen, H. N.; Pettersen, J.; Solli, C.; Hertwich, E. G. Investigating the Carbon Foot-print of a University - The case of NTNU. Journal of Cleaner Production 2013, 48, 39–47.

(17) Ozawa-Meida, L.; Brockway, P.; Letten, K.; Davies, J.; Fleming, P. Measuring Carbon Performance in a UK University through a Consumption-Based Carbon Footprint: De Montfort University Case Study. Journal of Cleaner Production 2013, 56, 185–198.

(18) Helmers, E.; Chang, C. C.; Dauwels, J. Carbon Footprinting of Universities World-wide: Part I—Objective Comparison by Standardized Metrics. Environmental Sciences Europe 2021, 33, 30.

(19) Achten, W. M. J.; Almeida, J.; Muys, B. Carbon footprint of science: More than flying. Ecological Indicators 2013, 34, 352–355.

(20) van Ewijk, S.; Hoekman, P. Emission Reduction Potentials for Academic Conference Travel. Journal of Industrial Ecology 2021, 25, 778–788.

(21) Stevens, A. R. H.; Bellstedt, S.; Elahi, P. J.; Murphy, M. T. The imperative to reduce carbon emissions in astronomy. Nature Astronomy 2020, 4, 843–851.

(22) Knödlseder, J.; Brau-Nogúe, S.; Coriat, M.; Garnier, P.; Hughes, A.; Martin, P.; Tibaldo, L. Estimate of the carbon footprint of astronomical research infrastructures. Nature Astronomy 2022, 6, 503–513.

(23) WRI, The Greenhouse Gas Protocol - A Corporate Accounting and Reporting Standard.; Report, 2015.

(24) Robinson, O. J.; Tewkesbury, A.; Kemp, S.; Williams, I. D. Towards a universal carbon footprint standard: A case study of carbon management at universities. Journal of Cleaner Production 2018, 172, 4435–4455.

(25) Robinson, O. J.; Kemp, S.; Williams, I. D. In HANDBOOK OF THEORY AND PRACTICE OF SUSTAINABLE DEVELOPMENT IN HIGHER EDUCATION, *VOL* 2 ; Filho, W. L., Skanavis, C., DoPaco, A., Rogers, J., Kuznetsova, O., Castro, P., Eds.; 2017; pp 441–452.

(26) Harangozo, G.; Szigeti, C. Corporate Carbon Footprint Analysis in Practice – With a Special Focus on Validity and Reliability Issues. Journal of Cleaner Production 2017, 167, 1177–1183.

(27) Gömez, N.; Cadarso, M.-Á.; Monsalve, F. Carbon Footprint of a University in a Multi-regional Model: The Case of the University of Castilla-La Mancha. Journal of Cleaner Production 2016, 138, 119–130.

(28) Mariette, J.; Blanchard, O.; Berné, O.; Aumont, O.; Carrey, J.; Ligozat, A.; Lellouch, E.; Roche, P.-E.; Guennebaud, G.; Thanwerdas, J.; Bardou, P.; Salin, G.; Maigne, E.; Servan, S.; Ben-Ari, T. An open-source tool to assess the carbon foot-print of research. Environmental Research: Infrastructure and Sustainability 2022, 2, 035008.

(29) Minx, J. C., et al. INPUT–OUTPUT ANALYSIS AND CARBON FOOTPRINTING: AN OVERVIEW OF APPLICATIONS. Economic Systems Research 2009, 21, 187– 216.

(30) Tukker, A.; Jansen, B. Environmental Impacts of Products: A Detailed Review of Studies. Journal of Industrial Ecology 2006, 10, 159–182.

(31) Suh, S. CEDA v.4.8 2014. https://vitalmetrics.com/environmental-databases.

(32) Ingwersen, W.; Li., M. Supply Chain Greenhouse Gas Emission Factors for US Industries and Commodities. Supply Chain Factors Dataset v1.0. https://doi.org/10.23719/1517796.

(33) Yang, Y.; Ingwersen, W. W.; Hawkins, T. R.; Srocka, M.; Meyer, D. E. USEEIO: A new and transparent United States environmentally-extended input-output model. Journal of Cleaner Production 2017, 158, 308–318.

(34) ADEME, Documentation générale : Ratio monétaires. https://bilans-ges.ademe.fr/fr/accueil/documentation-gene/index/siGras/1.

(35) Rebitzer, G.; Ekvall, T.; Frischknecht, R.; Hunkeler, D.; Norris, G.; Rydberg, T.; Schmidt, W. P.; Suh, S.; Weidema, B. P.; Pennington, D. W. Life cycle assessment: Part 1: Framework, goal and scope definition, inventory analysis, and applications. Environment International 2004, 30, 701–720.

(36) Carbon disclosure project. https://www.cdp.net.

(37) Majeau-Bettez, G.; Strømman, A. H.; Hertwich, E. G. Evaluation of Process- and Input–Output-based Life Cycle Inventory Data with Regard to Truncation and Aggregation Issues. Environmental Science & Technology 2011, 45, 10170–10177.

(38) GES 1point5. https://apps.labos1point5.org/ges-1point5.

(39) Labos 1point5. https://labos1point5.org.

(40) RTE, Bilan électrique 2019. https://assets.rte-france.com/prod/public/2020-06/bilan-electrique-2019_1_0.pdf.

(41) IEA, Global Energy & CO2 Status Report 2019. https://www.iea.org/reports/global-energy-co2-status-report-2019/emissions.

(42) OWID, Carbon intensity of electricity, 2022. https://ourworldindata.org/grapher/carbon-intensity-electricity?tab=tabledonn\’ee2021.

(43) Farley, M.; Nicolet, B. P. Re-use of labware reduces CO2 equivalent footprint and running costs in laboratories. bioRxiv 2022, 2022.01.14.476337.

(44) Herth, A.; Blok, K. Quantifying universities’ direct and indirect carbon emissions – the case of Delft University of Technology. International Journal of Sustainability in Higher Education 2023, 24, 21–52.

(45) Stadler, K. et al. EXIOBASE 3: Developing a Time Series of Detailed Environmentally Extended Multi-Regional Input-Output Tables. Journal of Industrial Ecology 2018, 22, 502–515.

(46) ThermoFisher, IsoFootprint: paving the way to sustainable isotope analysis; White paper, 2021.

(47) Raccary, B.; Loubet, P.; Peres, C.; Sonnemann, G. Applying Life Cycle Assessment to an analytical method: a case study on GC-MS analysis of pesticides in freshwater. submitted 2023,

(48) Font Vivanco, D. The role of services and capital in footprint modelling. The International Journal of Life Cycle Assessment 2020, 25, 280–293.

(49) PŔe sustainability, Environmental impacts of a satellite mission. https://pre-sustainability.com/customer-cases/environmental-impacts-satellite-mission/.

(50) Wiedenhofer, D.; Smetschka, B.; Akenji, L.; Jalas, M.; Haberl, H. Household time use, carbon footprints, and urban form: a review of the potential contributions of everyday living to the 1.5C climate target. Current Opinion in Environmental Sustainability 2018, 30, 7–17.

(51) AMUE, La nomenclature d’achats NACRES. https://www.amue.fr/presentation/articles/article/la-nomenclature-dachats-nacres-nouvelle-version/, 2021.

