## Supplementary information for "Purchases dominate the carbon footprint of research laboratories"

**4 Comparison with Larsen et al 2012 carbon itensities 31**

**SI References 32**

### 1 Supplementary methods

NACRES codes are composed of two letters and two numbers. The first letter provides the general category of the purchase, the second letter designs the domain, the first number the sub-domain and the last number the type.

#### 1.1 Detailed macro approach

We used version 4.8 of the CEDA database, from 2014, with EFs calculated according to the Life Cycle Inventory method, expressed in kg$CO_2$e/USD of 2002 in retail price. They were converted to 2019 USD from detailed tables of inflation by sector between 2002 and 2011 and by taking the average inflation(*1*) for the US economy between 2011 and 2019 (13.7%). For year 2019, USD were then converted to € with the rate(*2*) of 1.12USD for 1€.

USEEIO EFs were gathered from *Supply Chain GHG Emission Factors for US Commodities and Industries* v1.0 (`https://doi.org/10.23719/1517796`). We used *Supply Chain Emission Factors with Margins*, and more specifically 2016_Summary_Commodity data, which take into account "direct and indirect GHG emissions associated with production of commodity [...] from cradle to the point of sale in kg per 2018 USD of that commodity or industry in purchaser price of that commodity or industry in the US". Note that these EFs are slightly different than EFs taken directly from the USEEIO model that do not include margins (https://www.epa.gov/land-research/us-environmentally-extended-input-output-useeio-t The Supply Chain USEEIO database is expressed in 2018 USD that were also converted to 2019 € using the same protocol. The CEDA and USEEIO databases provide the EFs associated with different GHGs directly in kg$CO_2$e. We therefore added these values to obtain the total EF. The year of calculation of the EFs in the ADEME database is not specified. We thus took these factors as they were, without making any correction for inflation.

Note that the ADEME database supposes production using French technology while the other two rely on American technologies, and therefore on emissions associated with these technologies. Our method mixes data from these three databases and therefore assumes a single, French-American production technology. Taking into account different technologies by country have been shown to result in just a 15% difference in the average carbon footprint of a whole university(*3*), which is negligible compared to the uncertainties associated with the EEIO method (around 50%).

#### 1.2 Detailed meso approach

Based on corporate annual reports, we first identified companies which are mainly manufacturing products intended for research or similar analytical activities. We then selected the 15 companies that disclosed reasonably complete Scope 3 emissions, ie companies that published reliable emissions of the Purchased goods and services category (Table S4). GHG emission data were obtained either from the Carbon Disclosure Project (CDP) database (https://www.cdp.net), or from corporate reports. A limitation of this approach is that, in November 2022, reasonably complete and reliable GHG emissions (including upstream scope 3) were available only for few large companies, listed in Tabs. S4 and S2. When companies did not report emissions for capital goods, a value corresponding to 3% of total emissions was hypothesized. Missing employee commuting emissions were calculated on the basis of 1.5 t$CO_2$e/employee. For each company, the emissions by category are given in supplementary table Tab. S2.

To compute the carbon intensity of companies, we added scope 1, scope 2, scope 3 upstream emissions and the category downstream transportation and distribution, as transportation to laboratories is most of the time included in the price of purchases. The relative part of downstream transportation is below few percents for most companies, with the exception of companies that deliver their products by air, for which the relative share can reach 24% of emissions. The carbon intensity of companies or meso monetary EFs ($EF_{meso}$) are then computed as $EF_{meso}$ = (scope 1+2+3 emissions)/(revenue). Financial data were obtained from corporate annual reports. In the case of pharmaceuticals, we used the monetary emission factor computed with a similar approach, on a large number of pharmaceutical companies, by the non-profit organization My Green Lab, and published in its 2021 report entitled "The carbon impact of Biotech & Pharma" (`https://www.mygreenlab.org/carbon\_impact\_of\_biotech\_and\_pharma.html`). We used 2019 data from the top 15 companies, with emissions from downstream scope 3 sector deduced (except transportation and distribution) to keep the same perimeter as the one used for other companies.

#### 1.3 Detailed micro approach

For 60 lab products distributed in 28 NACRES categories, we performed a single-impact cradle-to-gate life-cycle assessment. The choice of products was based on three criteria: 1) only mono-material products were considered (with the exception of LB broth); 2) importance in term of amount of purchases in at least one discipline, and 3) the secondary EFs needed to calculate the carbon footprint were already available in process-based life cycle inventory databases or in publications. LCA included raw material manufacturing, item manufacturing, transport to the local supplier and then transport to the laboratories (sterilization was not considered due to a lack of data). Both container (primary packaging) and content were considered. We based our analysis on references in use in the laboratory of one of the authors (Table S3). In the case of gas cylinders, two types of cylinder (a large one, L50, and a medium-size one, M20) were systematically considered. For each item, we used manufacturer information to determine the raw material composition. Each plastic or glass item was weighted using an OHAUS traveler balance. Secondary packagings (cardboard boxes for deliveries) were not included. For transport, we assumed that products were transported by sea and/or road directly from the place of production, determined using manufacturer information, to intermediate storage places of distributors and then to the final laboratory. Most of the EFs were obtained from the Ecoinvent database Version 3.8, using IPPCC 2013 GWP100 values of the geographical area of production and/or market-based values (Table S3). To account for GHG emissions not taken into account into LCA (Business travel, Employee commuting, Waste, Purchases of other goods and services and Non-attributable processes), the total $CO_2$e obtained was multiplied by 1.15 to obtain the micro EF of the product. This multiplication factor was determined based on emissions of plastic processing companies, whith limited research and development activities, and simple production lines. Results of carbon footprints are available in Table S3. Two carbon footprints were not derived from our calculations but from publications: disposable gloves(*4*), and a kit for the isolation and purification of nucleic acid (`https://www.qiagen.com/-/media/project/qiagen/qiagen-home/about-qiagen-website/documents/summary-life-cycle-assessment-qiagen-2020.pdf`). In this last case, emissions of Qiagen company not considered in the LCA such as business travel, waste, employees commuting, investments and part of scope 1 and 2 not attributable to production activities were added on the pro rata of a kit price to the company revenue.

The monetary carbon footprints were then computed by dividing the product carbon footprint by its price. Prices of items in NACRES categories H and N are 2019 prices from the French central purchasing office for research and education (UGAP, `https://www.ugap.fr/`), ensuring homogeneity of pricing among French laboratories. Unlike other items, prices of gas cylinders are not subject to a national tender and are negotiated by each laboratory in France. In our study, prices of gases are those of 2021 Air Liquide Industrie price offer to the Micalis laboratory. These prices were chosen after a comparison of prices of few gas references from two other laboratories, that showed that they are on the average. $EF_{micro}$ were then computed by averaging the monetary carbon footprints of products from the same NACRES category (Table S3).

#### 1.4 Constitution of the PER1p5 database

PER1p5 database was constructed by substituting some EFs of the NACRES-EF macro database by meso and micro EFs. $EF_{meso}$ of companies producing reasonably homogeneous types of goods (either instruments, consumables or services representing at least 80% of sales) were used: $EF_{meso}$ of Agilent, Bio-Rad and Thermo-Fisher were not considered as these companies cover too large business segments. An exception concerns Charles River, providing both services and laboratory animals, whose emission factor was used for laboratory animals. The 12 selected $EF_{meso}$ were attributed to 102 NACRES categories in the PER1p5 final database (Table S1). Similarly, not all computed $EF_{micro}$ are used in the PER1p5 database, but only those corresponding to relatively homogeneous NACRES categories. For inhomogeeous categories, we considered that $EF_{macro}$ computed with EEIOA approaches were preferable as the values of $EF_{micro}$ were too dependent on the choice of products for LCA. The relative standard deviation of monetary carbon footprints whithin a NACRES category was considered as a proxy for homogeneity: only $EF_{micro}$ of categories with relative standard deviation below 0.5 were used (Table S3). For gases, given the homogeneity of the NACRES categories (one category per type of gas and level of purity) and the impossibility to compute relative standard deviation (only two references per category), we replaced all $EF_{macro}$ with $EF_{micro}$. In two cases, a $EF_{micro}$ has been attributed to other NACRES categories than those of the products on which they were calculated: $EF_{micro}$ NB43 (reusable labware other than pipettes) has also been attributed to NB42 and NB44 categories and the $EF_{micro}$ NB74 has been attributed to NB71, NB72, NB73 and NB75 categories. 31 NACRES categories of the final PER1p5 database were attributed $EF_{micro}$ (see Table S1).

In addition to the EFs detailed above, we associated several tags to each NACRES type for data processing:

- The *method* tag indicates for a given NACRES wether macro, meso or micro EFS were used in the PER1p5 database.

- The *module* tag sets which module in GES 1point5 is used to calculate the emissions for a given NACRES type. It can take five different values: `purchases`, `vehicles`, `energy`, `travels`, `devices`. The monetary approach described here is only used to calculate the emissions of the NACRES types labeled `purchases`. For the other values, we used EFs based on physical flows as described in ref. *5*.

- The *category* tag classifies the purchases of the laboratory in 7 aggregated categories in order to identify action strategies. These categories are *lab.life* (Food, landscaping, leisure, building), *consumables* (Raw materials, chemicals/biologicals and living organisms), *lab.equipment* (Laboratory equipment and instruments), *transport* (professional travel, including lodging but excluding transport), *info* (audio-video equipment), *services* and *maintenance*.

#### 1.5 Uncertainties

We calculated two types of uncertainties in the PER1p5 database (Table S1):

- *per1p5.uncertainty.attr.kg.co2e.per.euro*: Attribution uncertainty of a macro EF calculated as the standard deviation from the three macro EFS: CEDA, USEEIO and ADEME. This uncertainty was not used throughout this work.

- *per1p5.uncertainty.80pct.kg.co2e.per.euro*: Calculated as 80% of the corresponding EF. This uncertainty was used for each EF throughout this work.

To calculated the uncertainty associated to a sum of emissions from different NACRES codes we proceeded in two steps. The tag *uncertainty.groupby* identify a group of EFs that *are not* independent from each other. Within this group uncertainties are summed linearly.

Outside this group EFs *are* considered independent from each other and thus variances are summed linearly.

#### 1.6 Perimeter of the GHG inventory

The perimeter includes all upstream and in-house laboratory emissions except those due to heavy investments (such as construction and large scientific infrastructures), waste, staff meals and some specific direct emissions such as anaesthetic gas or methane emitted by ruminants. It includes transport to the distributors (but not from the distributors to the laboratory). In particular each emission source in Fig. 4 in the Main Text includes:

- Purchases: all goods and services purchased by the lab except utilities, train and plane tickets and fuel (see Fig. S1 for details). It does include taxi and good transportation.
- Building energy: heat, electricity.
- Refrigeration gases.
- All professional travels.
- All commuting travels.
- Direct emissions by vehicles owned by le laboratory (cars, boats...).

#### 1.7 Data curation

Emissions and staff numbers were fed by volunteer laboratories into the GESpoint5 lab emissions database. We analysed only emissions sources labeled as *submitted* indicating that volunteer laboratories have checked their data. We statistically analysed the purchases emission per capita data to look for suspect outliers. The corresponding laboratories were contacted which confirmed the results and thus all data were kept in our analysis.

#### 1.8 Mitigation strategies

Seven mitigation strategies (MS) were investigated:

- MS1 assumes a 50% increase in the service life of laboratory equipments. The total emissions and the emissions from "lab.equipment" and of "repair and maintenance" categories were summed by domain. The footprint of equipments was divided by 1.5 and the footprint of repair and maintenance was multiplied by 1.5.

- MS2 assumes a pooling of 50% of laboratory equipments. For the pooling by discipline, the total emissions and the emissions of "lab.equipment" and of "repair and maintenance" were summed by discipline, while for the pooling at the regional scale, the total footprint and the footprint of "equipments" and of "repair and maintenance" were summed by administrative region if at least 9 GHG assessments were available (7 regions). The footprint of equipments was divided by 2 and the footprint of repair and maintenance was multiplied by 2. The results at the regional scale are the average of 7 regions.

- MS3 assumes an 80% decrease in the use of disposable plastic consumables (NACRES codes NB02, NB03, NB04, NB11, NB12, NB13, NB14, NB15, NB16 and NB17). It implies an 80% increase in the use of consumables for washing machines (NACRES code NB34). The first year, it also implies an increase in the purchases of glassware (NACRES code NB43; EF = $0.23 \pm 0.1$ kg $CO_2e$/€) for an amount equivalent of twice the amount of disposable plastic consumables. From the second year, a 5% breakage was assumed. The total footprint and the footprint of disposable plastic consumables and of consumables for washing machine were summed by domain.

- MS4 assumes a change in diet with an increase in the proportion of vegetarian menu for catering services paid by the laboratory. The emissions of catering services (NACRES codes AA63, AA64) were summed by domain. According to ADEME, the mean footprint of a traditional meal in France is 2.04 kg $CO_2e$ and the mean footprint of a vegetarian meal is 0.5 kg $CO_2$e. Assuming a 75 % conversion to vegetarianism, the footprint of catering services was divided by 3.

- MS5 assumes a 50% decrease in the purchases of furniture (NACRES codes AA43,AB02, AB03, AA52,AF01,CD43,CG11,CG12). Emissions of furniture were summed by discipline and divided by 2.

- MS6 assumes a 50% decrease in consumables. Emissions from "consumables" category were summed by discipline. The footprint of consumables was divided by 2.

- MS7 assumes a 50% decrease in IT purchases. Emissions from the devices module in GES1p5 and from 'info' category within the purchases module were summed by the domain and divided by 2.

Although monetary and aggregated approach that we have followed in this study does not allow evaluating mitigation strategies coming from choices of consumables or instruments with lower carbon footprint than their classical counterparts (supply-based strategies). Such mitigation strategies must be subject to specific estimates based on physical factors and data from suppliers. The difficulty of such mitigation strategies is that they require precise determination of the carbon footprints of one type of product from different manufacturers (or of different models of the same supplier). Few data exist for convenience goods that are part of lab purchases such as computers or printer toners. However, due to LCA limitations discussed above precise process-based carbon footprints are so far inexistent for laboratory equipments or specific consumables, limiting the possibility to evaluate mitigation strategies based on supplier specific processes for labs.

### 2 Supplementary Figures and Tables

#### 2.1 Emission factor tables

The three spreadsheets containing the emission factors for Tables S1-S3 are available at :

`https://dropsu.sorbonne-universite.fr/s/cN2BKerxR5EYxyM`.

Table S1: NACRES-EF database. Correspondence between NACRES codes, USEEIO codes, and macro, meso and micro EFs. Data for the CEDA database are not provided to respect the license. These data are available in the file `PER1p5_nacres_fe_database_v1-0-2023.xlsx`.

Table S2: Carbon emission and financial data from companies used to calculate meso EFs in Tab. S4. These data are available in the file `PER1p5_meso_data_full_v1-0-2023.xlsx`.

Table S 3: Detailed micro EFs. These data are available in the file `PER1p5_LCA_results_v1-0-2023.xlsx`.

Table S4: Meso EFs for different representative companies. Data calculated from *6*.

| **Types of sold goods** | **Company** | **Business segments** | **Carbon intensity (kg $CO_2$e/€)** |
|---|---|---|---|
| Instrumentation, consumables & services | Agilent | Laboratory instruments, consumables, chemicals & services | 0.13 |
| | Bio-Rad | Laboratory equipment, products and in-vitro diagnostic tests | 0.15 |
| | Thermo-Fisher | Laboratory instruments, consumables, chemicals & services | 0.38 |
| Equipment (>80% of sales) | Shimadzu | Analytical & measuring instruments and medical systems | 0.21 |
| | Hamamatsu | Optical devices | 0.35 |
| Consumables (> 80% of sales) | AbCam | Research antibodies and related products | 0.14 |
| | Ansell | Gloves and hygienic equipment | 0.74 |
| | Becton Dickinson | Medical delivery systems & culture media | 0.15 |
| | Illumina | New generation sequencing | 0.07 |
| | Qiagen | Molecular biology consumable kits | 0.16 |
| | Sartorius | Bioprocess products (filtration, separation etc.) | 0.15 |
| | Sigma-Aldrich (2015) | Chemicals for research | 0.45 |
| Consumables & services | Charles River | Drug development services and laboratory animals | 0.19 |
| Services | Eurofins | Analysis services, diagnostic tests and sequencing | 0.07 |
| | Wiley | Academic publishing and learning | 0.17 |

Tabe S5: List of the 28 large NACRES categories within which the 1431 NACRES codes using in this work are distributed.

| **NACRES cat. code** | **NACRES cat. label (french)** | **NACRES cat. label (english)** |
|---|---|---|
| A | APPROVISIONNEMENTS GENERAUX | GENERAL SUPPLIES |
| B | BATIMENTS - INFRASTRUCTURES - TRAVAUX - ESPACES VERTS | BUILDINGS - INFRASTRUCTURE - WORKS - GREEN SPACES |
| C | COMMUNICATION - CULTURE - DOCUMENTATION | COMMUNICATION - CULTURE - DOCUMENTATION |
| D | DEPLACEMENTS : TRANSPORT ET HEBERGEMENT DES PERSONNES | TRANSPORT AND ACCOMMODATION |
| E | ETUDES - CONSEILS - ASSURANCES - PI - RESSOURCES HUMAINES | STUDIES - CONSULTING - INSURANCE - PI - HUMAN RESOURCES |
| F | FRET / EXPEDITION / TRANSPORT / DEMENAGEMENT | FREIGHT / SHIPPING / TRANSPORT / MOVING |
| G | GAZ DE LABORATOIRE OU D'ATELIER - CRYOGENIE | LABORATORY OR WORKSHOP GASES - CRYOGENICS |
| H | HYGIENE ET SECURITE AU TRAVAIL | HYGIENE AND SAFETY AT WORK |
| I | INFORMATIQUE - TELECOMMUNICATIONS - AUDIOVISUEL | IT - TELECOMMUNICATIONS - AUDIOVISUAL |
| J | AMENAGEMENT DE LABORATOIRE ET DE SALLE DE TP | LABORATORY ROOM FITTINGS |
| K | EXPERIMENTATION ANIMALE - ELEVAGE ANIMAL - CONSERVATION ANIMALE - ETUDE DES ANIMAUX | ANIMAL EXPERIMENTATION - ANIMAL BREEDING - ANIMAL CONSERVATION - ANIMAL STUDIES |
| L | MEDICAL | MEDICAL |
| M | MICROSCOPIE - PROFILOMETRIE | MICROSCOPY - PROFILOMETRY |
| N | CHIMIE ET BIOLOGIE | CHEMISTRY AND BIOLOGY |
| O | OPTO - LASERS - MATERIEL D'OPTIQUE | OPTO - LASERS - OPTICAL EQUIPMENT |
| P | PHYSIQUE - PHYSIQUE NUCLEAIRE ET CORPUSCULAIRE | PHYSICS - NUCLEAR AND CORPUSCULAR PHYSICS |
| Q | EXPERIMENTATION VEGETALE | PLANT EXPERIMENTATION |
| R | ATELIER - MECANIQUE - AUTOMATIQUE | WORKSHOP - MECHANICS - AUTOMATION |
| S | SPECTROMETRIE - SPECTROSCOPIE - RAYONS X | SPECTROMETRY - SPECTROSCOPY - X-RAYS |
| T | ELECTRONIQUE / TEST, ENERGIE, MESURES | ELECTRONICS / TEST, ENERGY, MEASUREMENT |
| U | SCIENCES DE LA TERRE - GEOPHYSIQUE - ASTROPHYSIQUE - ARCHEOLOGIE | EARTH SCIENCES - GEOPHYSICS - ASTROPHYSICS - ARCHAEOLOGY |
| V | VIDE ET ULTRAVIDE : EQUIPEMENTS POUR LE VIDE ET POUR LES TECHNIQUES SOUS VIDE | VACUUM AND ULTRA-HIGH VACUUM : EQUIPMENT FOR VACUUM AND VACUUM TECHNOLOGY |
| W | NANOTECHNOLOGIES - MICROELECTRONIQUE | NANOTECHNOLOGIES - MICROELECTRONICS |
| X | DEPENSES HORS ACHATS | NON-PURCHASING EXPENSES |

#### 2.2 Data related to the section *Construction of the emission factor database* and Fig. 2 in the MT

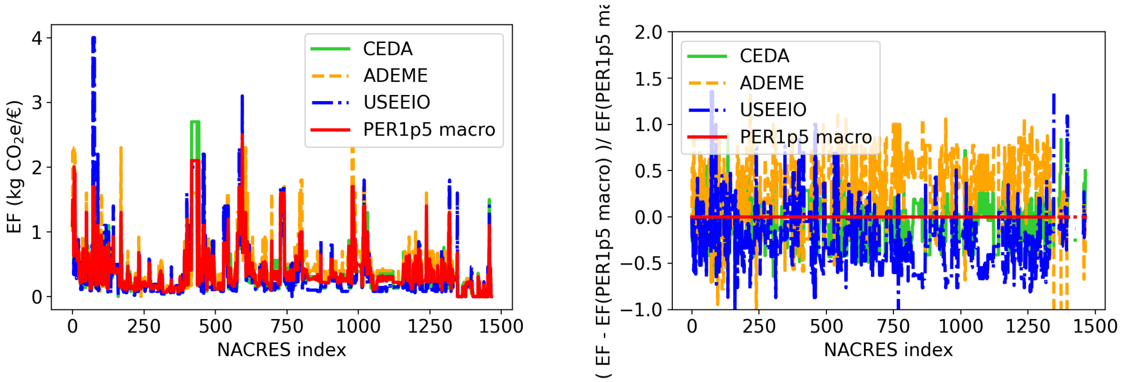

(a) (b)

Figure S1: Comparison of the 4 macro NACRES-EF databases for goods and services in the purchases module. (a) EF vs. NACRES index and (b) relative difference between the EF of a database and the EF of PER1p5 macro database.

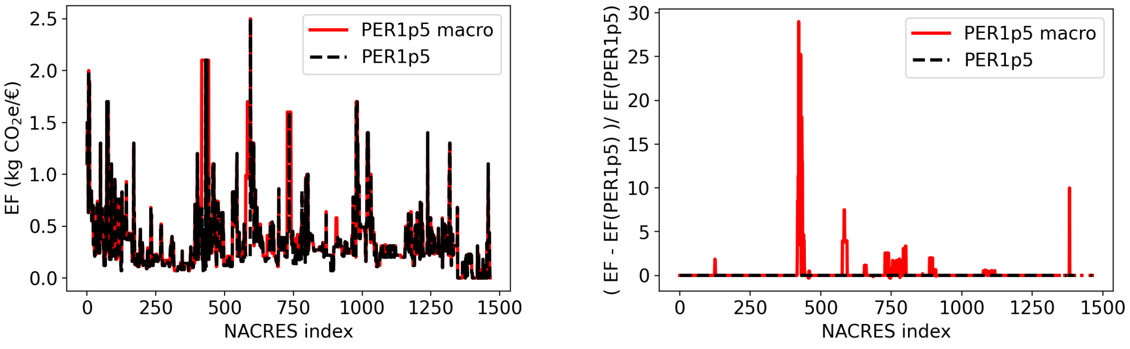

(a) (b)

Figure S2: Comparison of the PER1p5 macro and PER1p5 NACRES-EF databases for goods and services in the purchases module. (a) EF vs. NACRES index and (b) relative difference between the EF of a database and the EF of PER1p5 database.

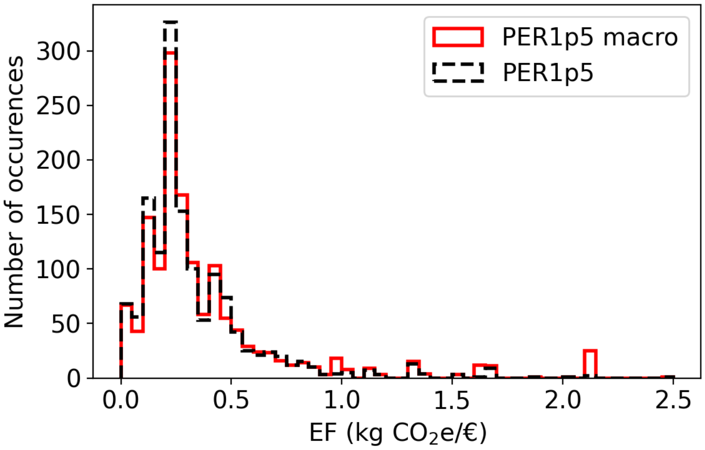

Figure S3: Histograms of the emission factors associated to the 1431 NACRES codes for the Purchases module for the database PER1p5 macro and its daughter PER1p5, where 138 emissions factors have been replaced by their meso/micro counterpart.

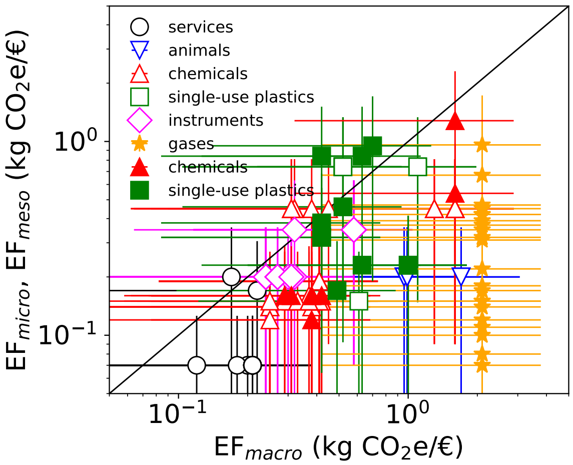

Figure S4: Meso (open symbols) and micro (filled symbols) emission factors vs. PER1p5 macro EF for different types of purchases, with error bars corresponding to a relative error of 80%. Corresponds to Fig. 2B in the MT but with error bars.

#### 2.3 Data related to the section *The distribution of carbon intensities in the laboratory research economy* and Fig. 3 in the MT

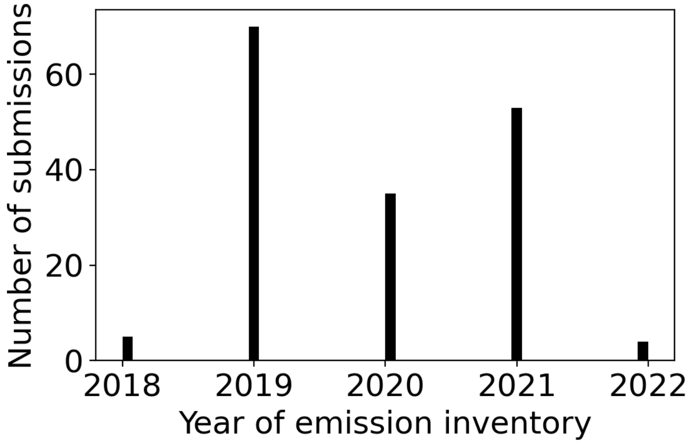

Figure S5: Histogram of submission year for all 167 purchases submissions from 105 laboratories.

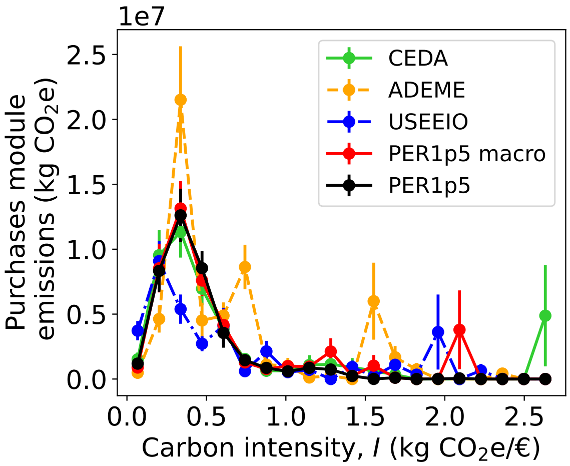

Figure S6: Distribution of carbon intensities within the GES 1point5 laboratory emission database calculated with all five NACRES-EF databases. For each bin in the $x$ axis, the corresponding carbon intensity $I$ is multiplied by the total amount of purchases in € to calculate the purchases emissions associated to that $I$. $n_s = 167$ GHG submissions averaged over $n_l = 108$ distinct laboratories.

Table S6: Statistics of the distribution of emission factors (EF) within each NACRES-EF database and within the GES 1point5 lab emission database for the five NACRES-EF databases used in this work. All the quantities are in kg $CO_2$e/€ and s.d. is the standard deviation.

| | **NACRES-EF database** | | | **GES 1point5 lab emission database** | | |
|---|---|---|---|---|---|---|
| **NACRES-EF database** | **Mean** | **Median** | **s.d.** | **Mean** | **Median** | **s.d.** |
| USEEIO | 0.33 | 0.18 | 0.45 | 0.29 | 0.28 | 0.09 |
| CEDA | 0.37 | 0.25 | 0.42 | 0.34 | 0.34 | 0.08 |
| ADEME | 0.47 | 0.40 | 0.41 | 0.43 | 0.44 | 0.10 |
| PER1p5 macro | 0.39 | 0.27 | 0.38 | 0.35 | 0.35 | 0.08 |
| PER1p5 | 0.33 | 0.24 | 0.28 | 0.31 | 0.30 | 0.07 |

Table S7: Total cumulated purchases module emissions from the 108 laboratories having submitted their inventories to the GES1point5 database, associated uncertainties and emissions relative to database PER1p5 per NACRES-EF database.

| **NACRES-EF database** | **tot. cum. em.** | **tot. uncertainty** | **rel. cum. em.** |
|---|---|---|---|
| CEDA | $4.5 \times 10^4$ | $0.5 \times 10^4$ | 1.15 |
| ADEME | $5.6 \times 10^4$ | $0.6 \times 10^4$ | 1.43 |
| USEEIO | $3.6 \times 10^4$ | $0.4 \times 10^4$ | 0.92 |
| PER1p5 macro | $4.6 \times 10^4$ | $0.5 \times 10^4$ | 1.16 |
| PER1p5 | $3.9 \times 10^4$ | $0.3 \times 10^4$ | 1.00 |

#### 2.4 Data related to the section *Purchases and electricity dominate laboratory emissions* and Fig. 4 in the MT

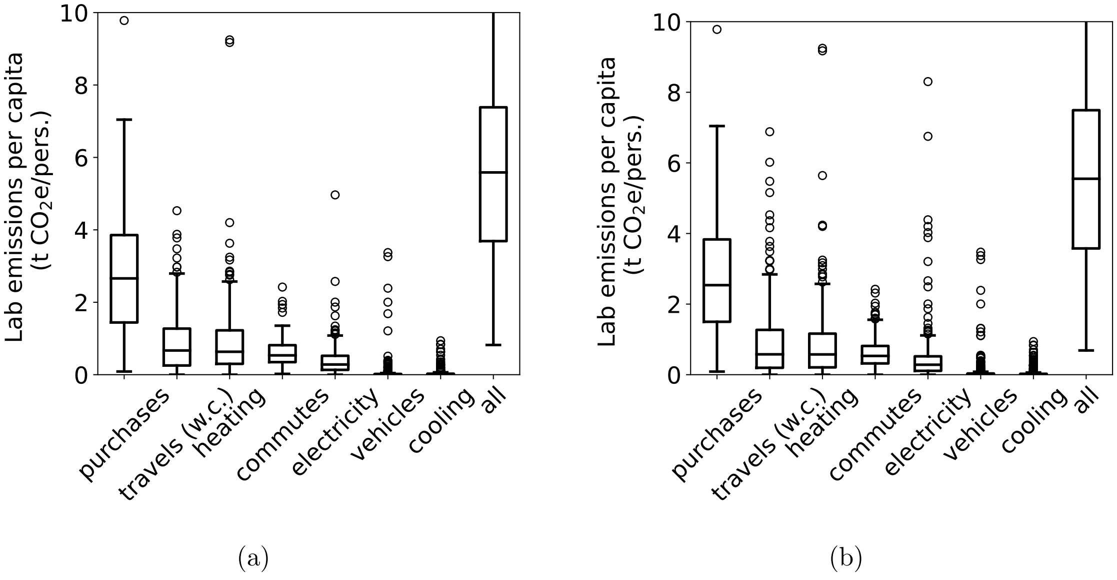

Figure S7: Comparison of laboratory emissions per capita per emission type when GHG submissions are averaged for each lab (a) or not (b). Panel (a) is identical to Figure 4A. Medians, boxes and bars are indistinguishable, there are only minor differences in outliers between the two panels.

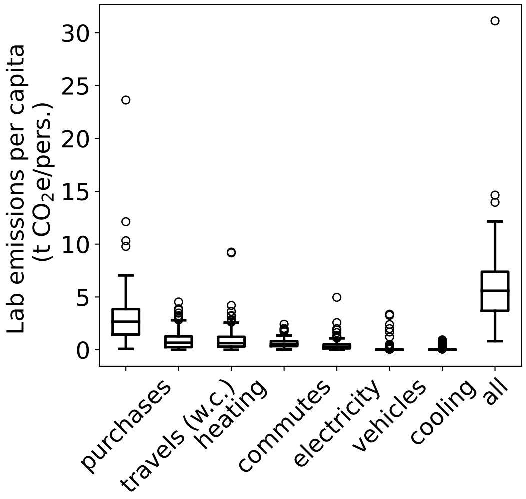

Figure S8: Boxplot of laboratory emissions per capita per emission source. $n_l \geq 190$ for all types except for purchases ($n_l = 105$). w.c. indicates that emissions associated to plane transportation were calculated with contrails(*5*). Same data as in Fig. 4 in the MT but without truncating the $y$ axis.

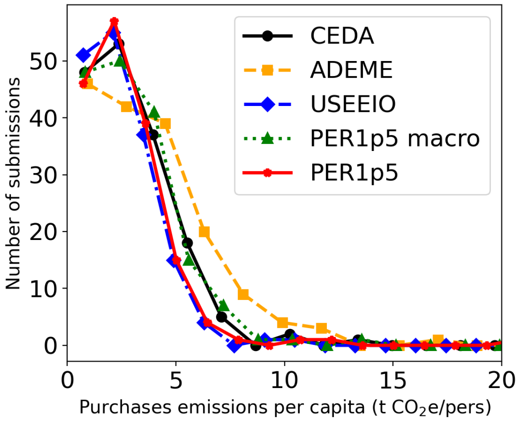

Figure S9: Distribution of purchases emissions within the BGES in the GES 1point5 laboratory emission database, depending on the NACRES-EF database used. Here, all 167 purchases submissions are considered.

Table S8: Statistics of laboratory emissions (top) and of GHG footprint submissions for each source considered. Staff is counted either by considering 1 professor = 0.5 (g1p5) or =1 (full). For each source the $n_s$ submissions are averaged per laboratory. Throughout this work we used the g1p5 way of counting staff for accounting the typical 50% time dedicated by professors in France to research activities

| **Source** | **Mean** (t $CO_2$ e/pers.) | **Relative man** (%) | **Median** (t $CO_2$ e/pers.) | **Quartiles** (t $CO_2$ e/pers.) |
|---|---|---|---|---|
| purchases | 3.2 | 51.6 | 2.7 | (1.4, 3.9) |
| travels (w.c.) | 0.9 | 14.5 | 0.7 | (0.3, 1.3) |
| heating | 0.9 | 14.5 | 0.6 | (0.2, 1.2) |
| commutes | 0.6 | 9.7 | 0.5 | (0.3, 0.8) |
| electricity | 0.5 | 8.1 | 0.3 | (0.1, 0.5) |
| vehicles | 0.1 | 1.6 | 0.0 | (0, 0) |
| cooling | 0.1 | 1.6 | 0.0 | (0, 0) |
| all | 6.2 | 100.0 | 5.6 | (3.7, 7.4) |

| **Source** | **# submissions** ($n_s$) | **# labs** ($n_l$) | **# staff (g1p5)** | **# staff (full)** |
|---|---|---|---|---|
| purchases | 162 | 105 | 20158.5 | 22764 |
| travels (w.c.) | 312 | 203 | 36464.5 | 41107 |
| heating | 289 | 190 | 33826.5 | 38128 |
| commutes | 288 | 197 | 33786 | 38027 |
| electricity | 289 | 190 | 33826.5 | 38128 |
| vehicles | 302 | 197 | 35384 | 39864 |
| cooling | 289 | 190 | 33826.5 | 38128 |
| all | 142 | 98 | 17269.5 | 19512 |
| purchases w/o devices | 162 | 108 | 21179 | 23891 |
| devices | 225 | 153 | 26712.5 | 30185 |

Table S9: Statistics of purchases module emissions (w/o devices) per capita within the 167 submissions, depending on the database used.

| **Database** | **Average em.** per cap (Ton $CO_2$e/pers.) | **Median em.** per cap (Ton $CO_2$e/pers.) | $\sigma$ (Ton $CO_2$e/pers.) |
|---|---:|---:|---:|
| USEEIO | 2.73 | 2.20 | 3.35 |
| CEDA | 3.32 | 2.72 | 3.82 |
| ADEME | 4.15 | 3.54 | 4.24 |
| PER1p5 macro | 3.40 | 2.78 | 3.78 |
| PER1p5 | 2.94 | 2.32 | 3.39 |

#### 2.5 Data related to the section *Purchases emissions are correlated to budget and research domain* and Fig. 5 in the MT

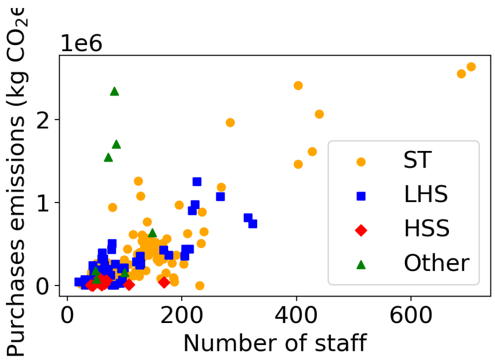

(a)

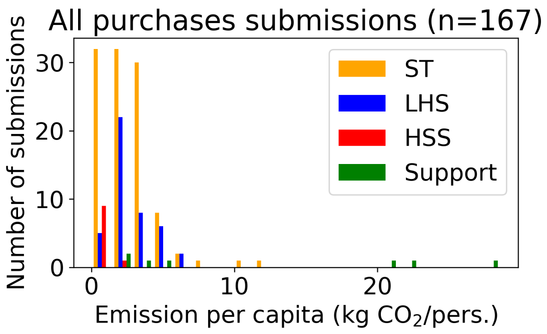

(b)

Figure S10: Purchases module emissions vs. number of staff (left) and corresponding histograms of purchase module emissions per capita (right). HSS: Human and social sciences, LHS: Life and health sciences, ST: Science and technology.

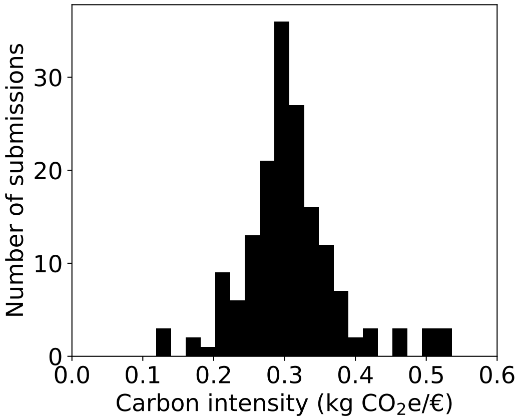

Figure S11: Histogram of purchases module carbon intensities for all 167 submissions from all domains.

Table S10: Definition of the research domains. In the GES 1point5 laboratory emission database each lab is assigned an HCERES code (hceres.code) identifying its subdiscipline. One lab may be attributed several subdisciplines. Here we have considered only the first indicated subdiscipline to identify a laboratory. For the domains we have preferred to use the MESRI (French Research and higher education ministry) notation instead of the HCERES one. The only difference is that the HCERES domain *Sciences de la vie et de l'environnement (SVE)* is recalled *Sciences du vivant et de la santé* (Life and health sciences, LHS). This last name represents better the subdisciplines included. `Hceres_disciplines_final.csv` at `https://dropsu.sorbonne-universite.fr/s/cN2BKerxR5EYxyM`.

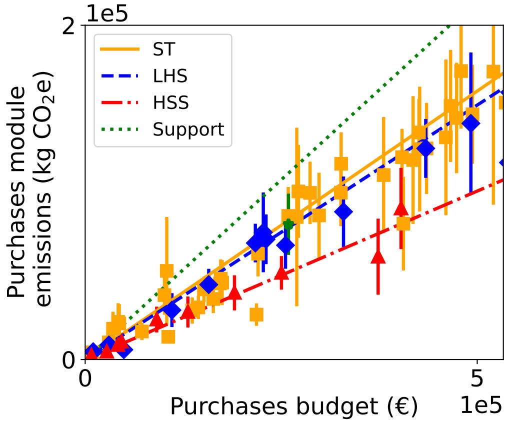

Figure S12: Zoom of Fig. 5A in the MT. Purchases module emissions vs. budget for all GHG laboratory footprints in the GES 1point5 lab emission database. Error bars corresponds to one standard deviation calculated as described in Methods. Lines are linear fits with zero intercept, whose results are provided in Tab. S11.

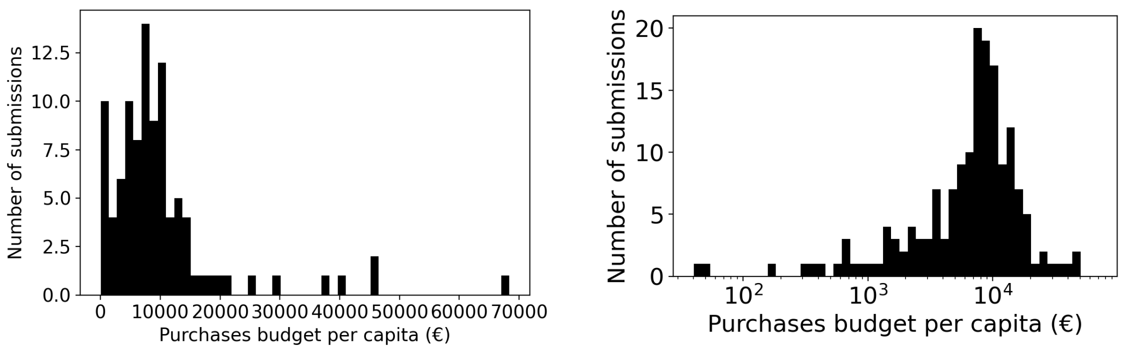

Figure S13: Histogram of purchases budget per capita with x-axis in lin (left) or log (right).

Table S11: Linear fits of purchases emissions vs. purchases budget for different domains in Fig. 5A (Slope and $R^2$) and average and standard deviation of carbon intensity from Fig. 5B. All values are in kg $CO_2$e/€ except for $R^2$.

| Domain | Slope | $R^2$ | $\bar{I}$ | $\sigma_I$ |
|---|---|---|---|---|
| Sciences and technology (ST) | 0.32 | 0.97 | 0.32 | 0.07 |
| Life and health sciences (LHS) | 0.30 | 0.97 | 0.29 | 0.05 |
| Human and social sciences (HSS) | 0.20 | 0.96 | 0.22 | 0.04 |
| Support | 0.43 | 0.96 | 0.36 | 0.10 |
| All | 0.33 | 0.96 | 0.31 | 0.07 |

#### 2.6 Data related to the section *The typology of purchases emissions depend on research domain* in the MT

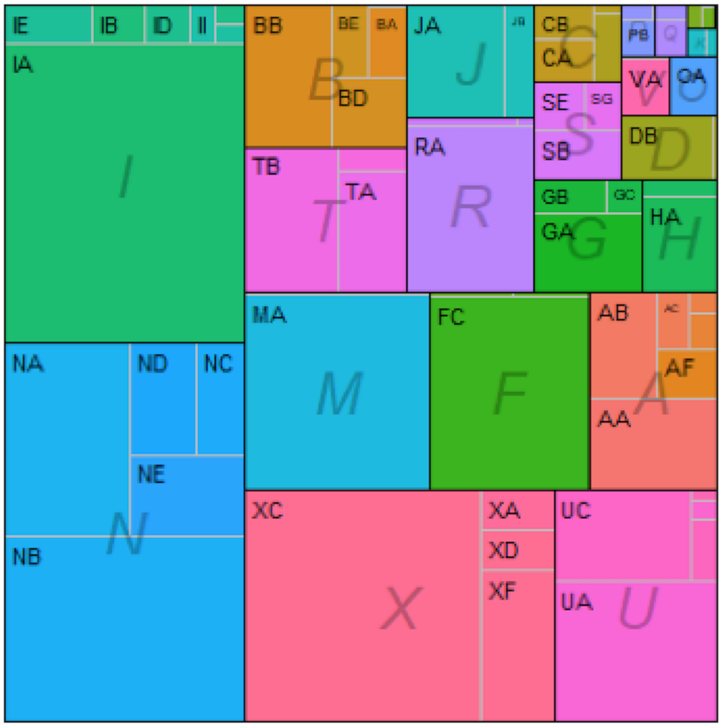

Figure S14: Treemap representation of the proportion of NACRES domains in the C footprint of purchases of a research laboratory in oceanic and atmospheric sciences.

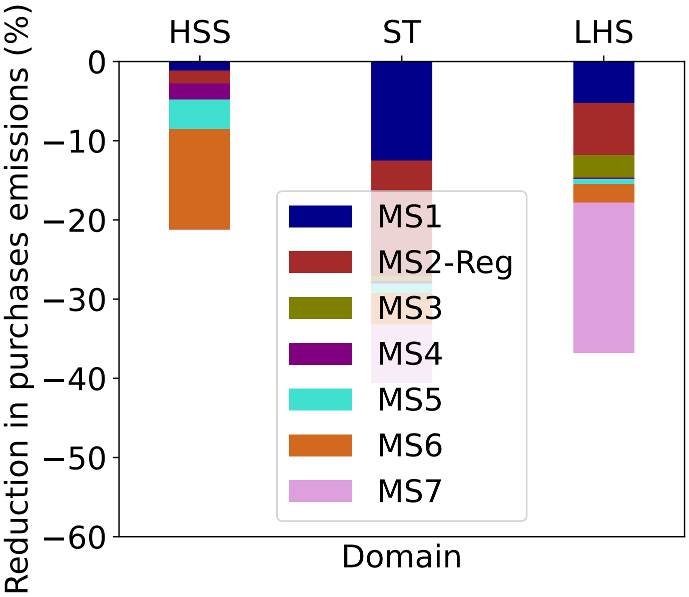

Figure S15: Relative reduction of the *purchases* laboratory emissions by research domain expected within the GES 1point5 lab emission database for the seven mitigation strategies considered. MS1: +50% of lab equipment life-time; MS2: 50% pooling of lab equipment by region (-Reg); MS3: replace 80% of plastic by glass; MS4: 75% vegetarian catering; MS5: −50% in furniture purchases; MS6: −50% in IT purchases; MS7: −50% in consumable purchases. $n_s = 135$ submissions corresponding to $n_l = 93$ laboratoires. ST: science and technology ($n_l = 64$), LHS: life and health sciences ($n_l = 23$), HSS: human and social sciences ($n_l = 6$) laboratories.

#### 2.7 Statistics on staff and laboratory domains in the GES 1point5 laboratory emission database

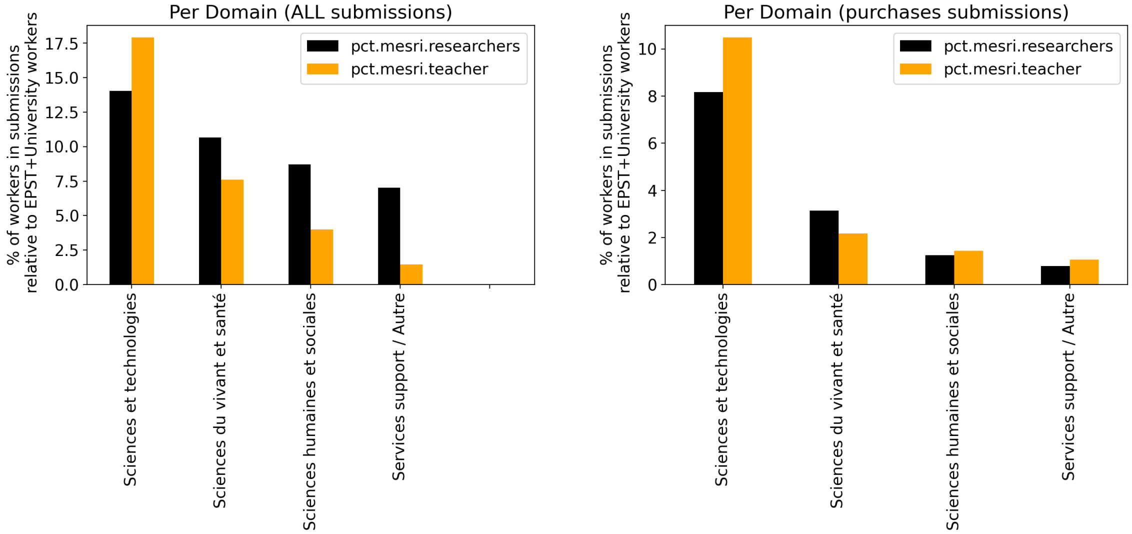

Figure S16: Statistics of the share of staff per research domain relative to tall the staff working in such domain within France for all submissions (left) and purchase submissions (right). This was done for two types of staff only: researchers and professors (called teacher in the legend). For each lab a weigthed sum between primary and secondary domain was considered (ponderation by 1 if only 1 domain and by 0.5 if 2). French statistics for te total number of researches and professors per domain were from refs. *7* and *8*.

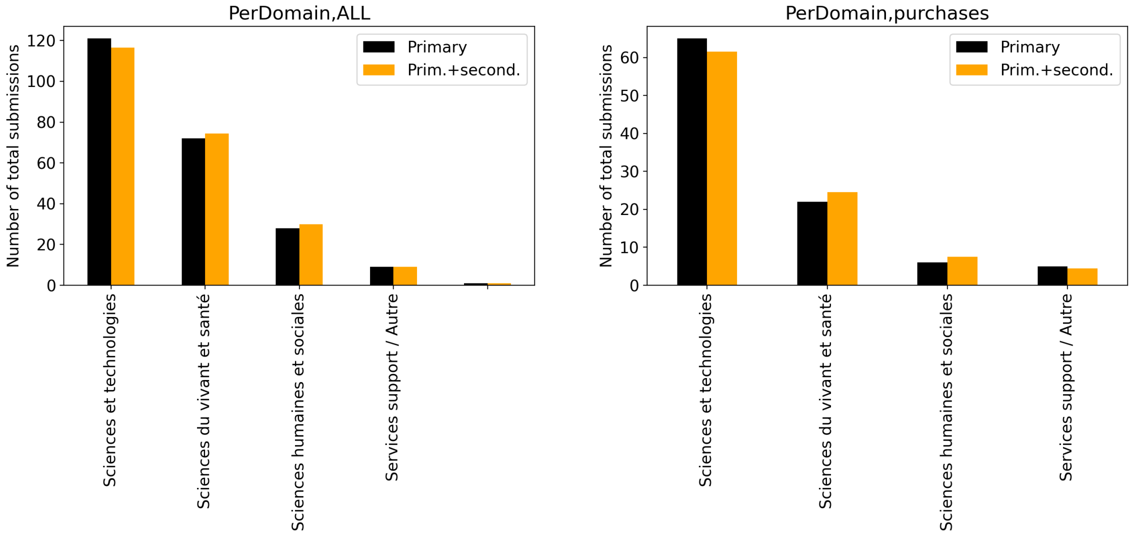

Figure S17: Statistics of number of submissions per research domain for all submissions (left) and purchase submissions (right). For each lab a weigthed sum between primary and secondary domain was considered (ponderation by 1 if only 1 domain and by 0.5 if 2).

### 3 Considerations about prices and technologies

#### 3.1 Inflation

EFs were corrected by inflation from their reference year to the year 2019 as detailed above. However, the purchases expenses reported by the laboratories in GES 1point5 laboratory emission database were not corrected from inflation because 94% of the inventories corresponded to the years 2019-2021 and the inflation in this period was < 4% in France. As in 2022 inflation reached < 9% for sectors outside energy, the webtool GES 1point5 will soon provide a correction to inflation for the EFs.

#### 3.2 Price parity

Tab. S12 provides a comparison of purchasing power parity between France and US for different sectors. The differences between the two countries < 10% for all sectors except Health and Miscellaneous goods and services. These small differences justify to mix the databases in US prices (CEDA, USEEIO) with the ones with French prices (ADEME and micro).

Table S12: Comparison of 2017 purchasing power parity (PPP) fro France, US and OECD countries in national currency per US dollar. Data from OECD (`https://stats.oecd.org/OECDStat_Metadata/ShowMetadata.ashx?Dataset=PPP2017&ShowOnWeb=true&Lang=en`)

| Country | Food and non-alcoholic beverages | Household furnishings, equipment and maintenance | Health | Restaurants and hotels | Miscellaneous goods and services | Machinery and equipment | Consumer goods | Non-durable goods | Semi-durable goods | Durable goods | Total services |
|---|---|---|---|---|---|---|---|---|---|---|---|
| France | 0.921 | 0.95 | 0.686 | 1.03 | 0.857 | 0.905 | 0.95 | 0.948 | 0.959 | 0.957 | 0.87 |
| U.S. | 0.988 | 1.01 | 1.27 | 1.01 | 1.11 | 0.975 | 0.959 | 0.938 | 1.01 | 0.977 | 1.21 |
| OECD | 1 | 1 | 1 | 1 | 1 | 1 | 1 | 1 | 1 | 1 | 1 |

#### 3.3 Production technologies

One may wonder wether the carbon intensity of the US and French economies are close enough such that one can average EEIO databases from each country. Fig. S18 compares the distribution of production EFs within the EXIOBASE 3 database for France and US. The differences are sufficiently small such that we can hypothesize that these two economies have similar carbon intensities for the types of goods considered in the PER1p5 database.

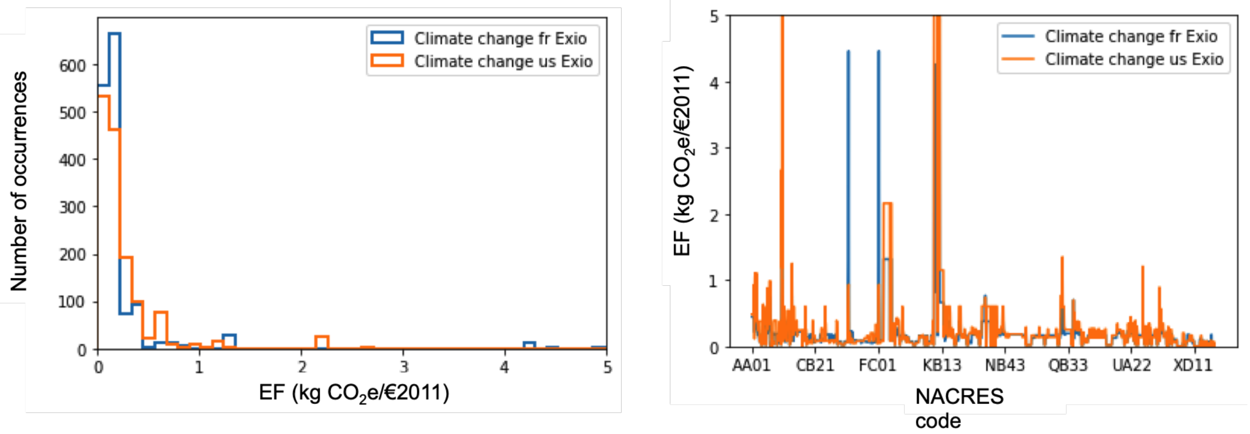

Figure S18: Comparison of EXIOBASE production emission factors within the NACRES-EF database between French (blue) and US production (orange). Note that these are production EFs and not consumption ones and thus they cannot be directly compared with Fig. 2A in the MT. Also here € values are given for year 2011. Data from EXIOBASE 3.8.2(*9*). We used values from matrix $M$ (MRIO extension multipliers (total requirement factors of consumption)), we converted EXIOBASE categories to USEEIO categories via the correspondence file and then automatically associated the EFs in $M$ by country to the NACRES code using Tab. S1.

### 4 Comparison with Larsen et al 2012 carbon itensities

Table S13: Purchases intensities for different faculties and domains from Larsen et al.(*10*). The figures in NOK for All and for each seven faculties come for the original work. Figures in € and for the three domains considered in this work were calculated from those data.

| Faculty / Domain | Emission intensity (kg $CO_2$e / (NOK 2009) ) | Emission intensity (kg $CO_2$e / (€ 2019) ) |
|---|---|---|
| All | 0.050 | 0.39 |
| Sciences and technology (ST) | 0.050 | 0.39 |
| Life and health sciences (LHS) | 0.046 | 0.36 |
| Human and social sciences (HSS) | 0.051 | 0.39 |
| Architecture | 0.039 | 0.30 |
| Humanities | 0.052 | 0.41 |
| Math and engineering | 0.051 | 0.40 |
| Engineering | 0.046 | 0.36 |
| Medecine | 0.041 | 0.32 |
| Natural Science | 0.052 | 0.41 |
| Social Science | 0.049 | 0.38 |

### SI References

1. US inflation calculator. https://www.usinflationcalculator.com/.

2. Frequency currency rates. https://freecurrencyrates.com/en/exchange-rate-history/EUR-USD/2019/cbr.

3. Gómez, N.; Cadarso, M.-Á.; Monsalve, F. Carbon Footprint of a University in a Multiregional Model: The Case of the University of Castilla-La Mancha. *Journal of Cleaner Production* **2016**, *138*, 119–130.

4. Rizan, C. A.-O.; Reed, M. A.-O.; Bhutta, M. F. Environmental impact of personal protective equipment distributed for use by health and social care services in England in the first six months of the COVID-19 pandemic. *Journal of the Royal Society of Medicine* *114*, 250–263.

5. Mariette, J.; Blanchard, O.; Berné, O.; Aumont, O.; Carrey, J.; Ligozat, A.; Lellouch, E.; Roche, P.-E.; Guennebaud, G.; Thanwerdas, J.; Bardou, P.; Salin, G.; Maigne, E.; Servan, S.; Ben-Ari, T. An open-source tool to assess the carbon footprint of research. *Environmental Research: Infrastructure and Sustainability* **2022**, *2*, 035008.

6. Carbon disclosure project. https://www.cdp.net.

7. SIES, *Note Flash du SIES. L'emploi scientifique dans les organismes de recherche en 2018*; Report, 2019.

8. Adedokun, F.; Tourbeaux, J. *Note de la DGRH - Enseignement supérieur - no 9 - Octobre 2022. Les personnels enseignants de l'enseignement supérieur du ministère de l'Enseignement supérieur et de la Recherche Année 2021*; Report, 2019.

9. Stadler, K. et al. EXIOBASE 3 (3.8.2) Data set. 2021; https://doi.org/10.5281/zenodo.5589597.

10. Larsen, H. N.; Pettersen, J.; Solli, C.; Hertwich, E. G. Investigating the Carbon Footprint of a University - The case of NTNU. *Journal of Cleaner Production* **2013**, *48*, 39–47.
